# eIF1 and eIF5 dynamically control translation start site fidelity

**DOI:** 10.1101/2024.07.10.602410

**Authors:** Rosslyn Grosely, Carlos Alvarado, Ivaylo P. Ivanov, Oliver B. Nicholson, Joseph D. Puglisi, Thomas E. Dever, Christopher P. Lapointe

**Author notes:** These authors contributed equally (Rosslyn Grosely, Carlos Alvarado). Corresponding authors: Thomas E. Dever and Christopher P. Lapointe.

## Abstract

Translation initiation defines the identity of a synthesized protein through selection of a translation start site on a messenger RNA. This process is essential to well-controlled protein synthesis, modulated by stress responses, and dysregulated in many human diseases. The eukaryotic initiation factors eIF1 and eIF5 interact with the initiator methionyl-tRNA_i_^Met^ on the 40S ribosomal subunit to coordinate start site selection. Here, using single-molecule analysis of in vitro reconstituted human initiation combined with translation assays in cells, we examine eIF1 and eIF5 function. During translation initiation on a panel of RNAs, we monitored both proteins directly and in real time using single-molecule fluorescence. As expected, eIF1 loaded onto mRNAs as a component of the 43S initiation complex. Rapid (∼ 2 s) eIF1 departure required a translation start site and was delayed by alternative start sites and a longer 5’ untranslated region (5’UTR). After its initial departure, eIF1 rapidly and transiently sampled initiation complexes, with more prolonged sampling events on alternative start sites. By contrast, eIF5 only transiently bound initiation complexes late in initiation immediately prior to association of eIF5B, which allowed joining of the 60S ribosomal subunit. eIF5 association required the presence of a translation start site and was inhibited and destabilized by alternative start sites. Using both knockdown and overexpression experiments in human cells, we validated that eIF1 and eIF5 have opposing roles during initiation. Collectively, our findings demonstrate how multiple eIF1 and eIF5 binding events control start-site selection fidelity throughout initiation, which is tuned in response to changes in the levels of both proteins.

## INTRODUCTION

Translation initiation canonically begins using an AUG codon at the start of the coding region of a messenger RNA (mRNA). The initiation machinery recognizes the start site with single-nucleotide precision, which maintains the proper reading frame and avoids synthesis of truncated and potentially toxic peptides. Yet, initiation also can proceed via start sites that differ by a single nucleotide (e.g., CUG, UUG, GUG)^1,2^. These alternative start sites typically yield less synthesized protein and have regulatory roles^3–5^. How the translation initiation machinery discriminates different potential start sites and the mechanistic basis for the decreased efficiency on alternative start sites remains unclear.

Translation initiation is directed by eukaryotic initiation factors (eIFs) and the initiator methionyl-tRNA_i_^Met^ (Met-tRNA_i_^Met^). The process starts when a 43S initiation complex – the small (40S) ribosomal subunit bound by eIF1, eIF1A, eIF3, and the eIF2–GTP–Met-tRNA_i_^Met^ ternary complex – loads onto the mRNA, which is facilitated by eIF4 proteins bound near the 5’-m^7^G cap^6–8^. While directionally moving 5’ to 3’ along the 5’UTR, the complex scans for translation start sites^9–11^, with selection enhanced by favorable Kozak sequence context^12^. A start site correctly placed at the ribosomal P site and base paired with the anticodon of Met-tRNA_i_^Met^ remodels the complex from a scanning-permissive conformation (P_scan_, aka P_out_) to a scanning-arrested conformation (P_in_)^9,10,13–15^. The remodeling coincides with eIF5-stimulated hydrolysis of GTP by eIF2^16,17^, which enables eIF2 departure. To complete initiation, a second GTPase, eIF5B, binds and collaborates with eIF1A to reposition Met-tRNA_i_^Met^ and allow the large (60S) ribosomal subunit to join and form the 80S ribosome^18–22^.

eIF1 and eIF5 play a central role during the unimolecular recognition of the start site by the anticodon of Met-tRNA_i_^Met^ on the initiation complex. In yeast, genetic and biochemical studies indicate that eIF1 discriminates ideal from suboptimal translation start sites; increased eIF1 levels suppress usage of alternative start sites, whereas decreased eIF1 levels or activity enhances their usage^23–31^. For eIF5, the opposite likely occurs; increased eIF5 activity enhances initiation on alternative sites^25,32^, perhaps by increasing the rate of eIF1 release from initiation complexes^33^. Similar effects may occur in mammalian systems^34–37^. These apparent concentration-dependent effects of eIF1 and eIF5 may be due to improper formation of 43S initiation complexes prior to their loading onto the mRNA^27,28,38–41^. However, the presence and role of eIF5 during formation of human 43S initiation complexes remains ambiguous.

Moreover, human initiation complexes that lack eIF1 improperly stall upstream of the start site, which can be rescued by subsequent addition of eIF1^42,43^. The concentration dependence of eIF1 and eIF5 effects therefore also could be explained by an alternate model where eIF1 and eIF5 dynamically bind and release initiation complexes to guide start site selection using distinct mechanisms.

eIF1 and eIF5 binding sites within initiation complexes overlap. Prior to recognition of the start site, eIF1 binds initiation complexes proximal to the ribosomal P site and the anti-codon stem and loop of Met-tRNA_i_^Met 44–53^. The presence of eIF1 restricts Met-tRNA_i_^Met^ to a scanning competent (P_scan_) conformation. Density for eIF1 disappears upon recognition of the translation start site, which correlates with movement of the 40S head region and Met-tRNA_i_^Met^ to the scanning-arrested (P_in_) conformation, partially occluding the eIF1 binding site^45,46,54–56^. By contrast, eIF5 is not observed in structures of initiation complexes prior to recognition of the start site^44–53^. Its absence could be due to structural flexibility or absence of the protein within the complex. Once stalled at the start site in the scanning arrested (P_in_) conformation, the N-terminal domain of eIF5 occupies the eIF1 binding site and directly contacts the anti-codon stem and loop of Met-tRNA_i_^Met 55,56^. However, when eIF1 and eIF5 bind and release initiation complexes, the nature of their exchange, and how their potential interplay might underpin recognition of the start site remain unknown.

To define how translation start sites are selected, we examined eIF1 and eIF5 directly during human translation initiation. We first developed single-molecule Förster resonance energy transfer (FRET) signals for each protein, which we integrated into our in vitro reconstituted human initiation system. Using equilibrium and real-time single-molecule spectroscopy assays, we determined the kinetics of eIF1 and eIF5 during initiation on a panel of mRNAs with various start sites. We reveal unexpected dynamics that explain prior cellular data from yeast, which we further demonstrated is conserved in human cells. Our findings provide a biophysical framework that indicates multiple kinetic steps underlie recognition of the translation start site.

## RESULTS

### An eIF1 fluorescence signal

To examine eIF1 function during translation initiation, we established a single-molecule FRET signal that monitors eIF1 on individual initiation complexes in vitro. Guided by published structures^44–53^, we purified and site-specifically labeled eIF1 with Cy5 fluorescent dye on Cys94 (**Fig. S1a**). In initiation complexes, this position is within ∼50 Å of the flexible U46 position on Met-tRNA_i_^Met^ (**Fig. 1a**). We therefore also conjugated Cy3 dye to U46 of synthetic tRNA_i_^Met^, which we aminoacylated with methionine to yield Met-tRNA_i_^Met^-Cy3 (hereafter tRNA_i_-Cy3) (**Fig. S1b**). We incubated tRNA_i_-Cy3 with purified eIF2 and a non-hydrolyzable GTP analog (GDPNP) to form the ternary complex, which we added to a mixture that contained purified 40S subunits, eIF1-Cy5, eIF1A, eIF3, eIF3j, eIF5, GDPNP, and ATP^22,57^. The resulting 43S initiation complex was incubated with a short (81 nt), 3’-biotinylated, unstructured model mRNA either with or without a translation start site (AUG) for 15 min at 37 °C. Complexes were tethered to neutravidin-coated imaging surfaces and visualized using total internal reflection fluorescence microscopy (TIRFm) upon excitation with a 532 nm laser at 23 °C (**Fig. S1c,d**).

**Figure 1.**
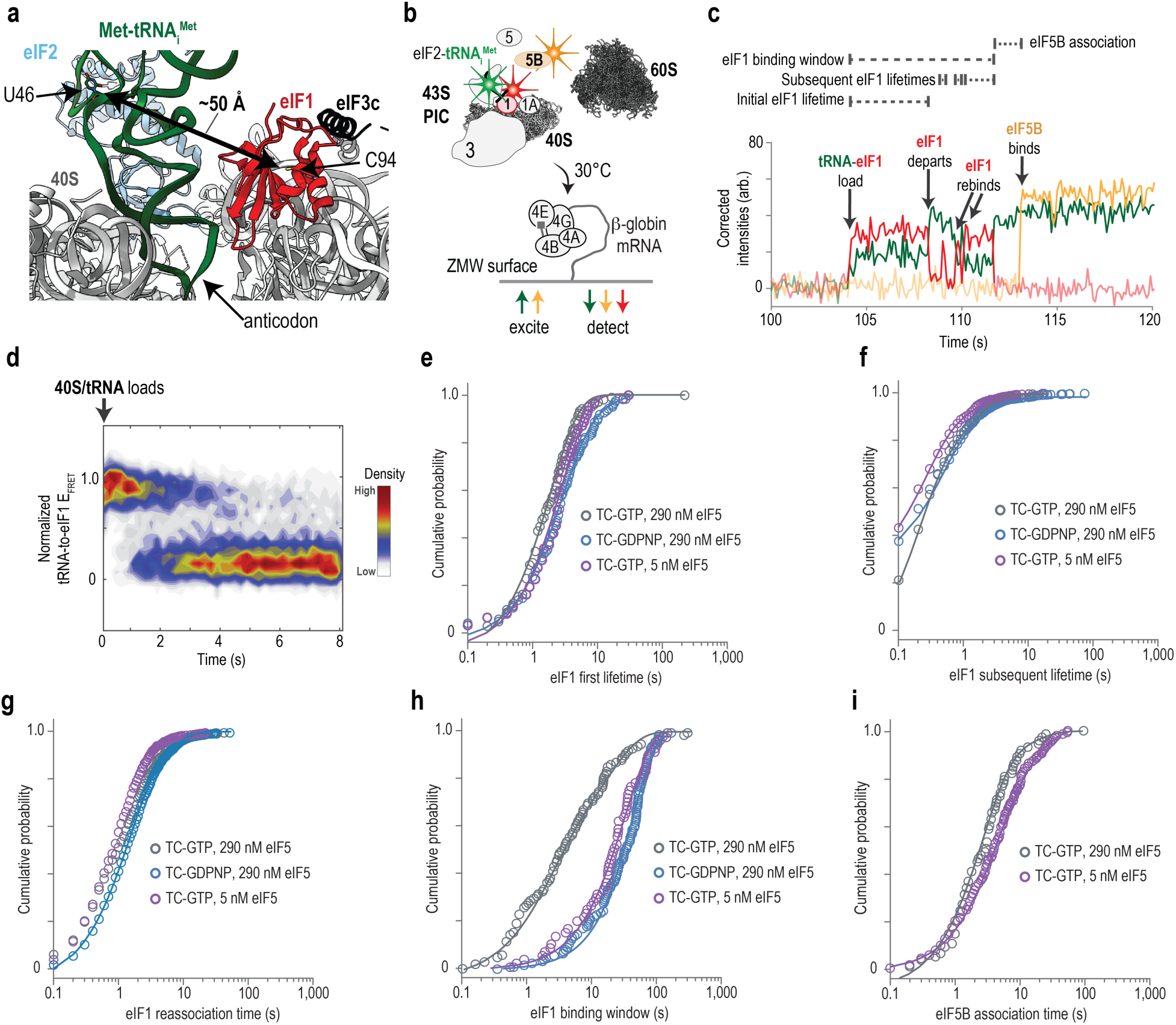
eIF1 stably binds and dynamically samples individual complexes. (**a**) Structure of a human translation initiation complex prior to recognition of the translation start site (PDB ID: 6ZMW). The positions of U46 on Met-tRNA_i_^Met^ (green) and C94 of eIF1 (red) are indicated, which are separated by about 50 Å. **(b)** Schematic of the real-time single-molecule assay. **(c)** Example single-molecule fluorescence data where tRNAi-Cy3 (green), eIF1-Cy5 (red), and eIF5B-Cy3.5 (orange) were monitored during translation initiation on β-globin mRNA. **(d)** Heat map of tRNA_i_-to-eIF1 FRET signal on all analyzed complexes synchronized to its initial appearance. **(e-i)** Cumulative probability plots of the indicated parameters (defined as in panel c and **Fig. S2a**). Lines represent fit to exponential functions, which we used to derive the reported rates. In all experiments here, eIF1-Cy5 was present at a final concentration of 40 nM (by Cy5) and unlabeled eIF5 was present at either 290 or 5 nM, as indicated. See **Table S1** for all rates and the number of complexes and binding events analyzed in each experiment.

eIF1 binds to distinct initiation complexes with variable kinetics at equilibrium. As we predicted given their proximity in structures, we observed pulses of eIF1-Cy5 fluorescent signal (red, FRET acceptor) and concomitant decreases in tRNA_i_-Cy3 intensity (green, FRET donor) when initiation complexes were formed on the model mRNA without a translation start site (**Fig. S1e**). We interpret tRNA_i_-to-eIF1 FRET as binding of eIF1 to the initiation complex and define the duration of the FRET signal as the eIF1 lifetime. The very high (0.9 ± 0.1) but broad FRET efficiency (E_FRET_) distribution suggests that the complex is structurally flexible at a rapid timescale. eIF1 binding also depended on the mRNA sequence: the median eIF1 lifetime on the initiation complex decreased by at least 7-fold (from 0.7 s to 0.1 s) on the mRNA with a translation start site (AUG) (**Fig. S1f,g**).

### eIF1 stably binds and dynamically samples individual complexes

We monitored eIF1 continuously throughout initiation using a prior real-time, single-molecule assay^22^ and our tRNA_i_-to-eIF1 FRET signal. We tethered full-length human β-globin mRNA with a 5’-m^7^G cap to an imaging surface within zero-mode waveguides (ZMWs), which facilitate four-color fluorescence detection in real time (**Fig. 1b**)^58^. Saturating concentrations of eIF4 proteins (4A, 4B, 4E, 4G), ATP, and GTP were then added to the tethered mRNA (see Methods). Separately, the 43S initiation complex with tRNA_i_-Cy3 (green) and eIF1-Cy5 (red) was preassembled as above, except using eIF2-GTP to allow initiation to proceed to completion. Upon start of data acquisition at 30 °C, a mixture with 10 nM 43S complex (by tRNA_i_-Cy3), 40 nM eIF5B-Cy3.5 (yellow), and 150 nM 60S subunits was added to the imaging surface. The samples were excited solely with a 532 nm laser, and the final eIF1-Cy5 and unlabeled eIF5 concentrations were 40 and 290 nM, respectively. We included eIF5B-Cy3.5 to demarcate initiation complexes that successfully progressed to late initiation steps (**Fig. 1b**)^22^. We report derived rates from the experiments throughout the text, and we aggregated all rates in a table (**Table S1**).

eIF1 bound stably and transiently sampled the same initiation complexes. As expected of a core 43S component, eIF1 loaded with the 43S initiation complex onto the mRNA, which was signaled by appearance of tRNA_i_-Cy3 (donor) to eIF1-Cy5 (acceptor) FRET (**Fig. S2a** and **Fig. 1c,d**). This initial eIF1 binding event persisted for about 2 s on average, which we define as the initial eIF1 lifetime (k_off, initial_ ≈ 0.5 ± 0.03 s^−1^) (**Fig. 1e**). Most initiation complexes (∼60%) contained multiple, subsequent eIF1 binding events (k_reassoc._ ≈ 0.55 ± 0.02 s^−1^ at 40 nM) that were about 10-fold briefer in duration (‘subsequent eIF1 lifetimes’, k_off, sub_. ≈ 5.5 ± 0.5 s^−1^) (**Fig. S2b** and **Fig. 1f,g**). The eIF1 binding window – defined as the dwell time from when eIF1 first loads onto the mRNA with the 43S complex until dissociation of the final eIF1 molecule – was 9 s in duration on average (at 40 nM eIF1) (**Fig. 1h**). Following final eIF1 departure, there was a 3 s delay prior to association of eIF5B (k_on, 5B_ ≈ 0.31 ± 0.02 s^−1^ at 40 nM) (**Fig. 1i**). Importantly, when measured relative to loading of the 43S initiation complex, the eIF5B association rate (0.1 ± 0.01 s^−1^ at 40 nM) matched the published rate obtained using unlabeled Met-tRNA_i_^Met^ and eIF1 (**Fig. S2c**)^22^. Thus, tRNA_i_-Cy3 and eIF1-Cy5 function identical to their unlabeled versions.

We examined how eIF1 kinetics vary in response to GTP hydrolysis by eIF2 in two ways. First, we replaced eIF2-GTP with eIF2-GDPNP and monitored eIF1-Cy5 during initiation. On these complexes, the duration of the initial eIF1 lifetime remained 2 s on average (k_off, initial_ ≈ 0.44 ± 0.05 s^−1^) (**Fig. 1e**). However, the complexes contained extensive subsequent eIF1 binding events (k_reassoc._ ≈ 0.57 ± 0.01 s^−1^ and k_off, sub._ ≈ 2.1 ± 0.1 s^−1^ at 40 nM) (**Fig. 1f,g** and **Fig. S2d**). The near continuous sampling by the protein extended the eIF1 binding window by at least 10-fold (from 9 s to >90 s) (**Fig. 1h**), which very likely was limited by photostability of the tRNA_i_-Cy3 signal. These complexes failed to progress to eIF5B association, consistent with prior work^18,22^. Second, we examined eIF1-Cy5 in the presence of eIF2-GTP and a limiting concentration of eIF5 (5 nM versus 290 nM), which is the GTPase activating protein for eIF2. Similar to eIF2-GDPNP, the initial eIF1 lifetime remained about 2 s in duration (k_off, initial_ = 0.34 ± 0.02 s^−1^), and the eIF1 binding window extended by at least 10-fold due to extensive eIF1 sampling (**Fig. 1e-h** and **Fig. S2e**). Thus, human translation initiation complexes contain multiple eIF1 binding events with distinct kinetics that respond to both GTP hydrolysis by eIF2 and reduced eIF5 levels.

### eIF1 dynamics depend on 5**’**UTR length and start site identity

To examine how eIF1 dynamics depend on the length of the 5’UTR, we generated two 5’ capped model mRNAs with either 50 nt or 200 nt long 5’UTRs comprised of CAA repeats. This design minimized potential RNA structures and enabled precise control of the translation start site identity and location without potential near cognate start codons (**Fig. 2a**). On both mRNAs, the canonical translation start site (AUG) was present in ideal Kozak context (**A**CC**<u>AUG</u>G**A). As expected, the 50 nt 5’UTR mRNA recapitulated eIF1 kinetics observed on β-globin mRNA (52 nt 5’UTR); at 40 nM eIF1-Cy5, the initial eIF1 lifetime (k_off, initial_ ≈ 0.36 ± 0.02 s^−1^), eIF1 reassociation rate (k_reassoc._ ≈ 1.3 ± 0.1 s^−1^), subsequent eIF1lifetime (k_off, sub._ ≈ 3.1 ± 0.3 s^−1^), eIF1 binding window (8 s), and eIF5B association rate (k_on, 5B_ ≈ 0.48 ± 0.05 s^−1^) essentially matched those on the native mRNA (**Fig. 2a**). Strikingly, the initial eIF1 lifetime lengthened by 2-fold on the 200 nt 5’UTR (from 2.7 s to 5.3 s, k_off, initial_ = 0.19 ± 0.02 s^−1^). eIF1 reassociation kinetics were unaffected. Consistently, the eIF1 binding window lengthened from 8 s to 11 s on the longer 5’UTR. Thus, the timing of the initial departure of eIF1 from initiation complexes depends on the length of the 5’UTR.

**Figure 2.**
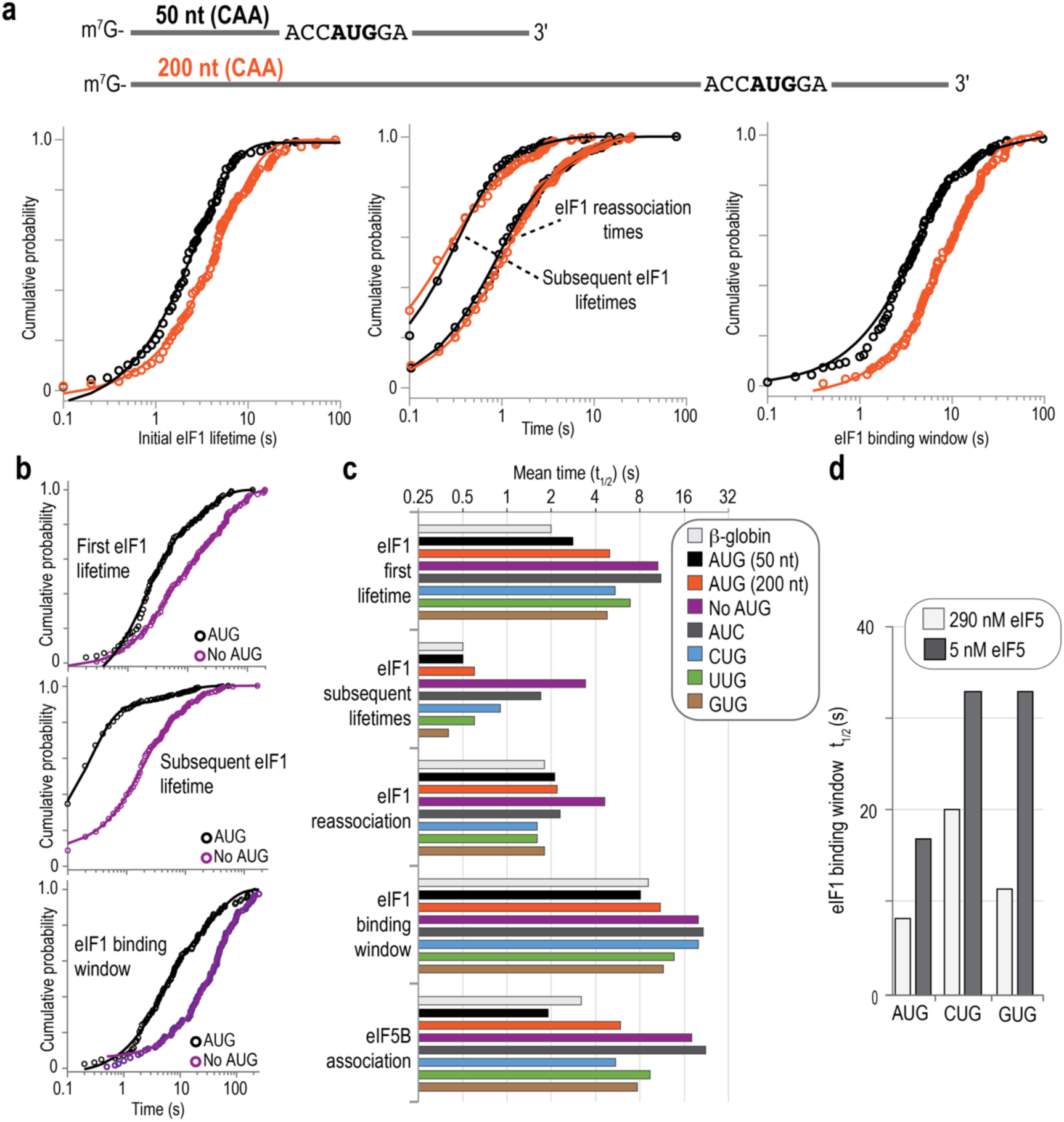
eIF1 kinetics depend on 5’UTR length and the identity of the translation start site. (**a**) Cartoon schematic of the model mRNAs with 50 nt (black) or 200 nt (orange) long 5’UTRs. Cumulative probability plots of the indicated eIF1 events observed on eIF5B-bound initiation complexes (successful). Lines represent fits to exponential functions. **(b)** Cumulative probability plots of the indicated eIF1 events observed on any loaded initiation complexes (both unsuccessful and successful). Lines represent fits to exponential functions. **(c)** Plot of the mean elapsed time (t_1/2_ value) of the indicated eIF1 parameters on the indicated mRNAs. All values plotted here were derived from events that occurred on eIF5B-bound initiation complexes (successful). **(d)** Plot of the mean elapsed time (t_1/2_ value) of the eIF1 binding window on the indicated mRNAs in the presence of 290 nM or 5 nM unlabeled eIF5. All values plotted here were derived from events that occurred on eIF5B-bound initiation complexes (successful). In all experiments, eIF1-Cy5 was present at a final concentration of 40 nM (by Cy5) and unlabeled eIF5 was present at either 290 or 5 nM, as indicated. See **Table S1** for all rates and the number of complexes and binding events analyzed in each experiment.

We hypothesized that eIF1 kinetics also would depend on the identity of the translation start site. To enable precise control, we used the 50 nt model 5’UTR from above and generated a panel that either lacked a start site (ACA) or contained CUG, UUG, GUG, or AUC alternative start sites. We examined eIF1 on each model mRNA and compared kinetics in two ways. First, we predicted that the success of initiation on mRNAs with different start sites would vary substantially (e.g., AUG vs. ACA). To capture those disparate outcomes, we examined eIF1 kinetics on all initiation complexes that loaded onto an mRNA, regardless of whether eIF5B associated (successful) or not (unsuccessful). Second, we hypothesized that eIF1 kinetics would vary for successful initiation complexes on the different start sites. We therefore examined eIF1 kinetics on the subset of initiation complexes that progressed to eIF5B association (our marker for success) on each model mRNA. In both scenarios, we further captured heterogeneity by comparing the mean elapsed times (*t*_1/2_ values), which proportionately integrated rates derived from either single-or double-exponential functions, as appropriate for each data set (for all rates, see **Table S1**).

Rapid initial eIF1 departure from initiation complexes required a translation start site. As expected, the initial eIF1 lifetime lengthened dramatically to 23 s on the model mRNA that lacked a translation start site (**Fig. 2b** and **Fig. S3a,b**). Similarly, the subsequent eIF1 lifetime lengthened to 4.7 s and the eIF1 reassociation time lengthened to 8.2 s. The collective effects lengthened the eIF1 binding window to more than 60 s. Nearly identical effects were observed on the model mRNA with an AUC start site, which is consistent with the well-established requirement for a G in the last position of the start codon for translation initiation (**Fig. S3a,b**). On both mRNAs, the extended eIF1 lifetimes likely were limited by photostability of the Cy5 dye and should be considered underestimates. On the subset of complexes that progressed to eIF5B association, the initial and subsequent eIF1 lifetimes lengthened by 5– and 6-fold (to 11 and 3.4 s) and delayed eIF5B association by 9-fold (to 18 s) (**Fig. 2c** and **Fig. S3c**). Such kinetics likely represent off-pathway initiation events, where initiation complexes settled for CAA start sites on these very short, artificial mRNAs. In stark contrast, eIF1 kinetics were identical on mRNAs with an AUG start site in poor Kozak context (**C**CC**<u>AUG</u>C**A) relative to the mRNA with the start site in ideal context (**Fig. S3c**).

Initiation on CUG, UUG and GUG-containing mRNAs led to lengthened and heterogeneous eIF1 kinetics. Compared to β-globin and the AUG model mRNA, the CUG, UUG, and GUG start sites increased by 2-3 fold the population of complexes that stalled with the initial eIF1 molecule stably bound (∼30 % of complexes) (**Figs. S3a,b** and **S3f-i**). These three alternative start sites also increased by 2-4 fold the subset of complexes that contained continuous eIF1 sampling events (∼6-18 % of complexes) (**Fig. S3f-i**). Both populations of complexes lacked eIF5B binding and likely represent stalled states. On complexes that progressed to eIF5B association and thus successfully progressed to downstream initiation steps, the initial and subsequent eIF1 lifetimes lengthened by 2.4-3.4 fold (to 5-7 s) and by 1-1.7 fold (to 0.4-0.9 s), respectively (**Fig. 2c** and **Fig. S3d**). eIF1 reassociation times were unaffected. The lengthened initial and subsequent eIF1 lifetimes lengthened the eIF1 binding window by 1.3-2 fold (to 11-20 s) relative to β-globin and the AUG model mRNA. The eIF5B association time also lengthened by 2.8-4.9 fold (to 5-9 s, at 40 nM eIF5B). Furthermore, the eIF1 binding window extended to more than 30 s when the eIF5 concentration was reduced from 290 nM to 5 nM on the CUG and GUG alternative start sites (**Fig. 2d** and **Fig. S3e**). Thus, the identity of the translation start site impacts the initial and subsequent eIF1 lifetimes and the eIF5B association rate.

### An eIF5 fluorescence signal

To monitor eIF5 during initiation, we leveraged available structures to label eIF5. In complexes that are stalled after recognition of the start site^55,56^, the N-terminal domain of eIF5 binds proximal to the anti-codon stem of Met-tRNA_i_^Met^ and the flexible U46 nucleotide (∼35 Å away) (**Fig. 3a**). We fused an 11 amino acid ybbR peptide tag to the N-terminus of eIF5, which we site-specifically labeled in vitro with Cy5 or Cy5.5 fluorescent dyes (eIF5-Cy5 and eIF5-Cy5.5) (**Fig. S4a**). This labeling position should yield FRET with tRNA_i_-Cy3. Given that yeast eIF5 can co-purify with eIF3^59^, we confirmed that our purified human eIF3 lacked eIF5 (and eIF1) using mass spectrometry (**Table S2**); thus, all single-molecule experiments below solely contained labeled eIF5-Cy5.5.

**Figure 3.**
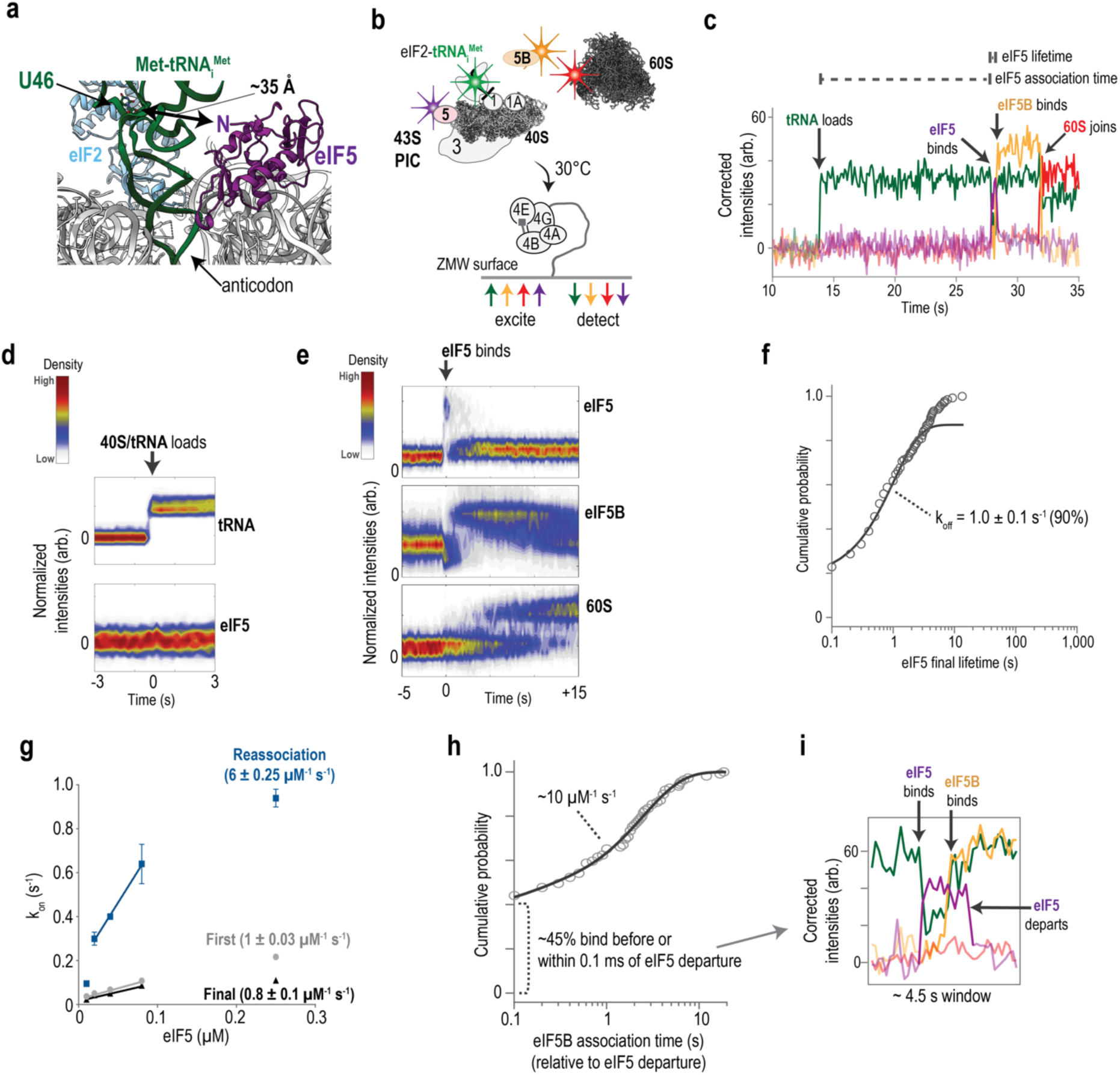
eIF5 transiently binds initiation complexes. (**a**) Structure of a human translation initiation complex present at the translation start site (PDB ID: 8OZ0). The positions of U46 on Met-tRNA_i_^Met^ (green) and the N-terminus of eIF5 (purple) are indicated, which are separated by about 35 Å. **(b)** Schematic of the real-time single-molecule assay. **(c)** Example single-molecule fluorescence data where tRNAi-Cy3 (green), eIF5-Cy5.5 (purple), eIF5B-Cy3.5 (orange), and 60S-Cy5 (red) were monitored during translation initiation on ϕ3-globin mRNA. **(d)** Heat map of the tRNA_i_-Cy3 (top) and eIF5-Cy5.5 (bottom) fluorescent signals on all analyzed complexes (the 40 nM eIF5 experiment) synchronized to initial appearance of the tRNA_i_-Cy3 signal. **(e)** Heat map of the eIF5-Cy5.5 (top), eIF5B-Cy3.5 (middle), and 60S-Cy5 (bottom) fluorescent signals on all analyzed complexes (the 40 nM eIF5 experiment) synchronized to appearance of the eIF5-Cy5.5 signal. **(f)** Cumulative probability plots of the observed eIF5 lifetime in the 40 nM experiment. The line represents the fit to a double-exponential function, which we used to derive the indicated rates. **(g)** Plot of the observed eIF5 association rates at the indicated concentrations. Lines represent fits via linear regression analysis to derive the indicated rates. **(h)** Cumulative probability plots of the observed eIF5B association time in the 40 nM experiment. The line represents a fit to an exponential function, which we used to estimate the indicated rate. **(i)** Example single-molecule data where the eIF5-Cy5.5 signal extended into the eIF5B-Cy3.5 binding event. In all experiments, the final concentration of eIF5-Cy5.5 is as indicated (by Cy5.5 dye) and unlabeled eIF1 was present at 290 nM. See **Table S3** for all rates and the number of complexes and binding events analyzed in each experiment.

We first examined eIF5-Cy5 on initiation complexes that contained eIF2-GDPNP and were equilibrated on the short model mRNAs with or without a translation start site. Upon excitation with a 532 nm laser using TIRFm, we observed pulses of eIF5-Cy5 fluorescent signal (purple, FRET acceptor) and concomitant decreases in tRNA_i_-Cy3 intensity (green, FRET donor) when initiation complexes were formed on the model mRNA with an AUG start site in ideal Kozak context (**Fig. S4b-d**). We interpret tRNA_i_-to-eIF5 FRET as binding of eIF5 to the initiation complex. The predominate observed E_FRET_ was very high (0.94 ± 0.09), with a minor and broad shoulder (0.72 ± 0.3) (**Fig. S4d**). eIF5 bound most initiation complexes (∼65%), and the protein remained bound for only hundreds of milliseconds on average (eIF5 lifetime, k_off1, eIF5_ = 3.4 ± 0.2 s^−1^) (**Fig. S4e**). By contrast, eIF5 binding was not detected on initiation complexes equilibrated on the mRNA without a translation start site (**Fig. S4f**). Thus, eIF5 binds dynamically to initiation complexes artificially stalled (by eIF2-GDPNP) at translation start sites.

### eIF5 transiently binds initiation complexes

We next monitored eIF5 directly during translation initiation in real time. We again tethered β-globin mRNA to an imaging surface within ZMWs and incubated the mRNA with saturating concentrations of eIF4 proteins (4A, 4B, 4E, 4G), ATP, and GTP (**Fig. 3b**). Separately, the 43S complex with tRNA_i_-Cy3 (green) and eIF5-Cy5.5 (purple) was preassembled, except using eIF2-GTP to allow progression of complexes to late initiation steps. Upon start of data acquisition at 30 °C, we added to the imaging surface a mixture that contained 10 nM 43S complex (by tRNA_i_-Cy3), 40 nM eIF5-Cy5.5, 40 nM eIF5B-Cy3.5 (yellow), and 150 nM 60S subunits labeled with Cy5 dye (red) on the C-terminus of uL18. Unlabeled eIF1 was present at 290 nM. During imaging, we excited the surface with 532 nm and 640 nm lasers simultaneously. eIF5-Cy5.5 occupancy thus was indicated by appearance of Cy3-to-Cy5.5 FRET, and by direct excitation and emission of the Cy5.5 dye. This excitation strategy circumvented any potential structural flexibility of the protein. It also enabled direct examination of both eIF5B and 60S subunit joining events, the final steps of initiation. Importantly, eIF5B and 60S subunit kinetics (relative to 40S complex loading) with tRNA_i_-Cy3 and eIF5-Cy5.5 present in the reaction were identical to when the unlabeled versions were present (**Fig. S5**)^22^. Thus, fluorescently-labeled Met-tRNA_i_^Met^ and eIF5 function identically to their unlabeled versions.

The 43S initiation complex loaded onto β-globin mRNA prior to eIF5 association. Loading of the 43S complex was indicated by appearance of tRNA_i_-Cy3 fluorescence and occurred independent of eIF5-Cy5.5 signal at all tested eIF5 concentrations (10, 20, 40, 80, and 250 nM) (**Fig. 3c,d**, **Fig. S6a,** and **Table S3**). Following a delay of 9 s after 43S loading (at 80 nM), 70-80% of initiation complexes contained a single, transient burst of eIF5-Cy5.5 signal that universally corresponded to a decrease in tRNA_i_-Cy3 intensity (k_off_ = 1.0 ± 0.1 s^−1^) (**Fig. 3c,e,f** and **Fig. S6b**). Given our excitation strategy, this observation indicates that eIF5 associated in a conformation that yields FRET with tRNA_i_-Cy3, rather than bind distally and rearrange at later time points. Furthermore, the limited concentration dependence of eIF5 association (0.8 ± 0.1 µM^−1^ s^−1^) suggests that an upstream slow step limits eIF5 association (**Fig. 3g** and **Fig. S6b**). On the 20-30% of complexes with multiple eIF5 binding events (**Fig. S6c,d**), eIF5 reassociated 6-fold more rapidly (5 ± 0.25 µM^−1^ s^−1^) (**Fig. 3g**), which suggests such complexes progressed past the limiting step.

eIF5 gates subsequent late steps in initiation. On 17-36% of initiation complexes, the eIF5 signal overlapped with appearance of the eIF5B-Cy3.5 signal (**Fig. 3h,i** and **Fig. S6e**). This finding demonstrates that both proteins can occupy initiation complexes simultaneously. On another 9-28% of complexes, loss of eIF5 signal was followed within 100 ms by eIF5B association, which is the temporal resolution of our experiments. Thus, about half of eIF5B association events (38-49%) occur just prior to or simultaneously with eIF5 departure from the initiation complex. For the remainder of complexes, eIF5B rapidly associated after eIF5 departure (0.41 ± 0.09 s^−1^ at 40 nM) (**Fig. 3h** and **Fig. S6b**). Given the incomplete labeling of eIF5 with dye (estimated at ∼50% labeled) and rapid eIF5 reassociation rate (5 ± 0.25 µM^−1^ s^−1^), we suspect that many of these slower eIF5B association events occurred on complexes that contained an unlabeled eIF5 protein. Once eIF5B bound, the 60S subunit joined and eIF5B departed the 80S initiation complex at rates identical to previous work (**Fig. S6b**)^22^. By contrast, eIF5 continuously sampled initiation complexes and failed to progress to eIF5B association when eIF2-GDPNP replaced eIF2-GTP in the reaction, as expected (**Fig. S6d**). Collectively, these findings strongly suggest that eIF5 binding allows eIF5B association. The two proteins can co-occupy initiation complexes for brief periods (< 1 s), but any potential direct interactions between the proteins occur transiently during initiation (k_off_ > 10 s^−1^)^60^.

### The identity of the translation start site alters eIF5 dynamics

We next examined the effects of different translation start sites on eIF5 dynamics using our panel of model RNAs. As expected, eIF5 bound 80% of initiation complexes that loaded onto the model mRNA with an AUG start site in either ideal or poor Kozak context (**Fig. 4a** and **Fig. S7a**). eIF5 association rates were similar (k_on_ = 0.13 ± 0.01 s^−1^ at 40 nM), whereas the eIF5 lifetime decreased by 2-fold to 0.4 s on the mRNA with a start site in poor context (k_off_ = 2.4 ± 0.2 s^−1^ versus 1.2 ± 0.07 s^−1^ in ideal context) (**Fig. S7b**). Thus, the context of the start site impacts eIF5 kinetics. The identity of the translation start site (in ideal contexts) also impacted binding of eIF5 to initiation complexes. The observed eIF5 binding efficiency decreased by about 1.5-fold on mRNAs where the A nucleotide was substituted for pyrimidines (CUG, UUG) (**Fig. 4a**). Substitution of either the A nucleotide for guanine (GUG) or the G nucleotide for cytosine (AUC) decreased eIF5 binding efficiency by 4-fold to baseline levels, as defined by the mRNA that lacked a start site (ACA). Modest increases in the number of eIF5 binding events also were observed on complexes loaded onto mRNAs with alternative start sites (**Fig. S7c**).

**Figure 4.**
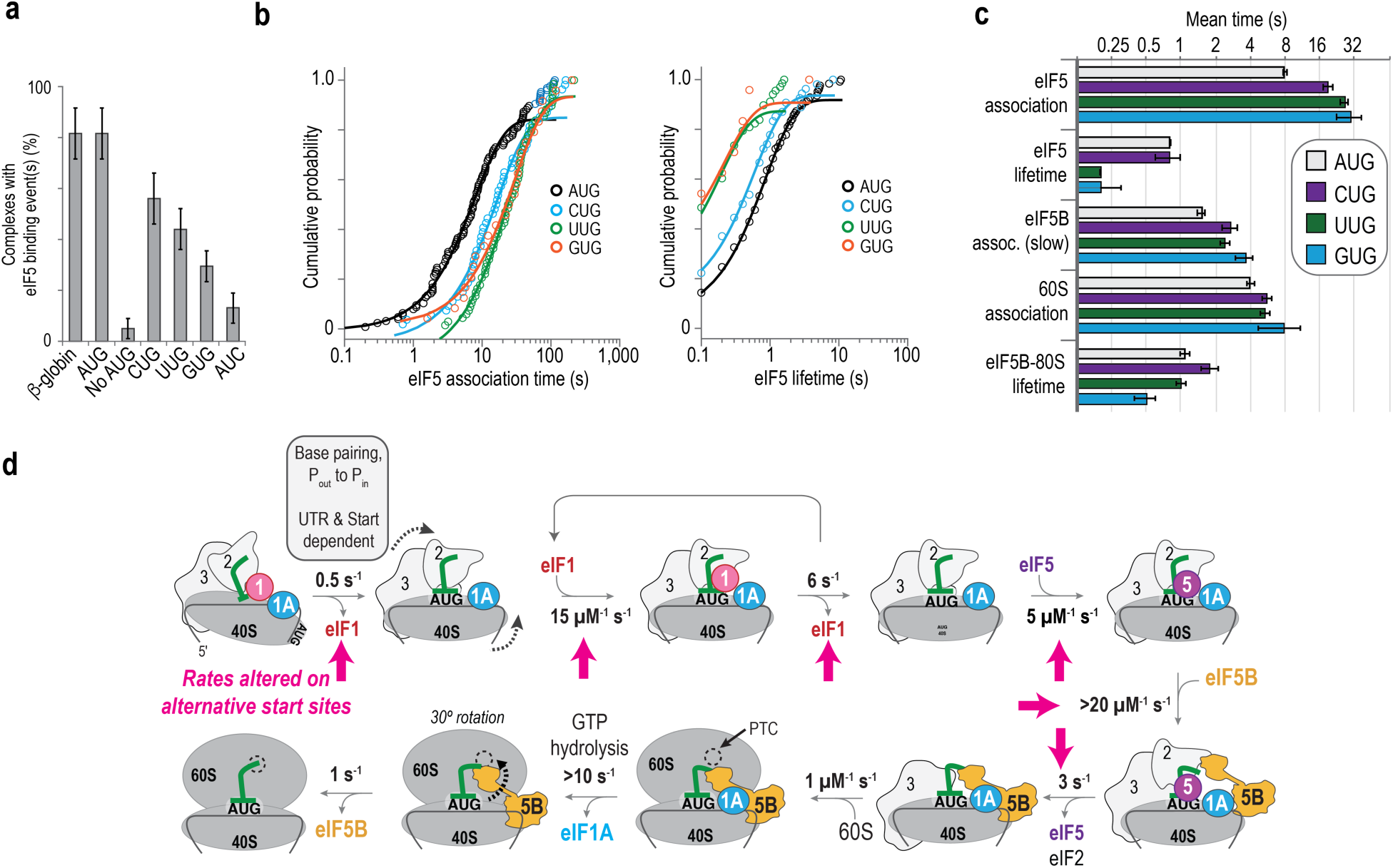
The identity of the translation start site alters eIF5 dynamics. (**a**) Plot of the percent of initiation complexes that contained at least one eIF5 binding event on the indicated mRNA. The percentage was corrected to account for an eIF5-Cy5.5 labeling efficiency of 50%. **(b)** Cumulative probability plots of the eIF5 association times (left) and eIF5 lifetimes (right) on the indicated mRNAs. For complexes with multiple eIF5 binding events, only the final eIF5 binding event was included in these plots. Lines represent fits to exponential functions. **(c)** Plot of the mean time (s) for the indicated kinetic parameters, which is defined as the reciprocal of the observed rate. The parameters are defined as in **Figure S6a. (d)** Kinetic model derived from our single-molecule studies on eIF1 and eIF5 (this study) and our previous study on eIF1A and eIF5B^22^. In all experiments, the final concentration of eIF5-Cy5.5 was 40 nM (by Cy5.5 dye) and unlabeled eIF1 was present at 290 nM. See **Table S3** for all rates and the number of complexes and binding events analyzed in each experiment.

While the alternative start sites largely abrogated eIF5 binding to initiation complexes, we also estimated kinetic effects by analyzing the rare subsets of complexes that did progress to eIF5 binding. The eIF5 association rate (at 40 nM) decreased by 2-, 3-, and 4-fold on the mRNAs with CUG (0.053 ± 0.01 s^−1^), UUG (0.038 ± 0.01 s^−1^), and GUG (0.034 ± 0.01 s^−1^) start sites relative to the AUG site in ideal context (0.13 ± 0.01 s^−1^) (**Fig. 4b,c** and **Fig. S7d**). The decreased rates lengthened the delay from loading of the 43S initiation complex until eIF5 association to nearly 30 s (from 8 s). Furthermore, the eIF5 dissociation rate increased by 4-fold on the mRNAs with UUG (5.1 ± 1 s^−1^) and GUG (4.6 ± 10 s^−1^) start sites relative to the CUG and AUG sites (both ∼1.2 ± 0.1 s^−1^), which decreased the eIF5 lifetime to 0.2 s. Thus, in the minority of cases where binding was observed, eIF5 associated more slowly with and departed more rapidly from initiation complexes on mRNAs with alternative start sites. These collective effects delayed eIF5B association. The number of initiation complexes where eIF5B associated during or immediately after (within 0.1 s) eIF5 binding decreased from ∼60% on the AUG mRNA to 23-37% on the alternative start sites (**Fig. S7e**). This effect mirrored the 2-3 fold decreased eIF5B slow association rates on the alternative start sites (**Fig. 4c** and **Fig. S7d**). Once initiation complexes progressed to eIF5B association, though, downstream initiation events on mRNAs with alternative start sites proceeded at rates essentially identical to the mRNA with an AUG start site (**Fig. 4c** and **Fig. S7d**).

### eIF1 and eIF5 have opposing roles in cells

Based on the kinetic model we derived from our single-molecule experiments (**Fig. 4d**) and prior studies examining overexpression of eIF1 and eIF5^36,37^, we hypothesized that changes in the relative abundance of eIF1 and eIF5 would impact translation in human cells. We therefore examined how both knockdown and overexpression of eIF1 and eIF5 impacted translation of mRNAs with ideal and alternative translation start sites. To enable precise control of the start site, we fused our panel of unstructured model 5’UTRs comprised of CAA repeats to the firefly luciferase coding region (**Fig. 5a**). We also examined two versions of the native eIF5 5’UTR, which contains two inhibitory uORFs with AUGs in poor context. In one, we maintained the uORFs, and in a second (used as a control), we eliminated the upstream AUGs, leaving solely the AUG at the start of the luciferase coding region.

**Figure 5.**
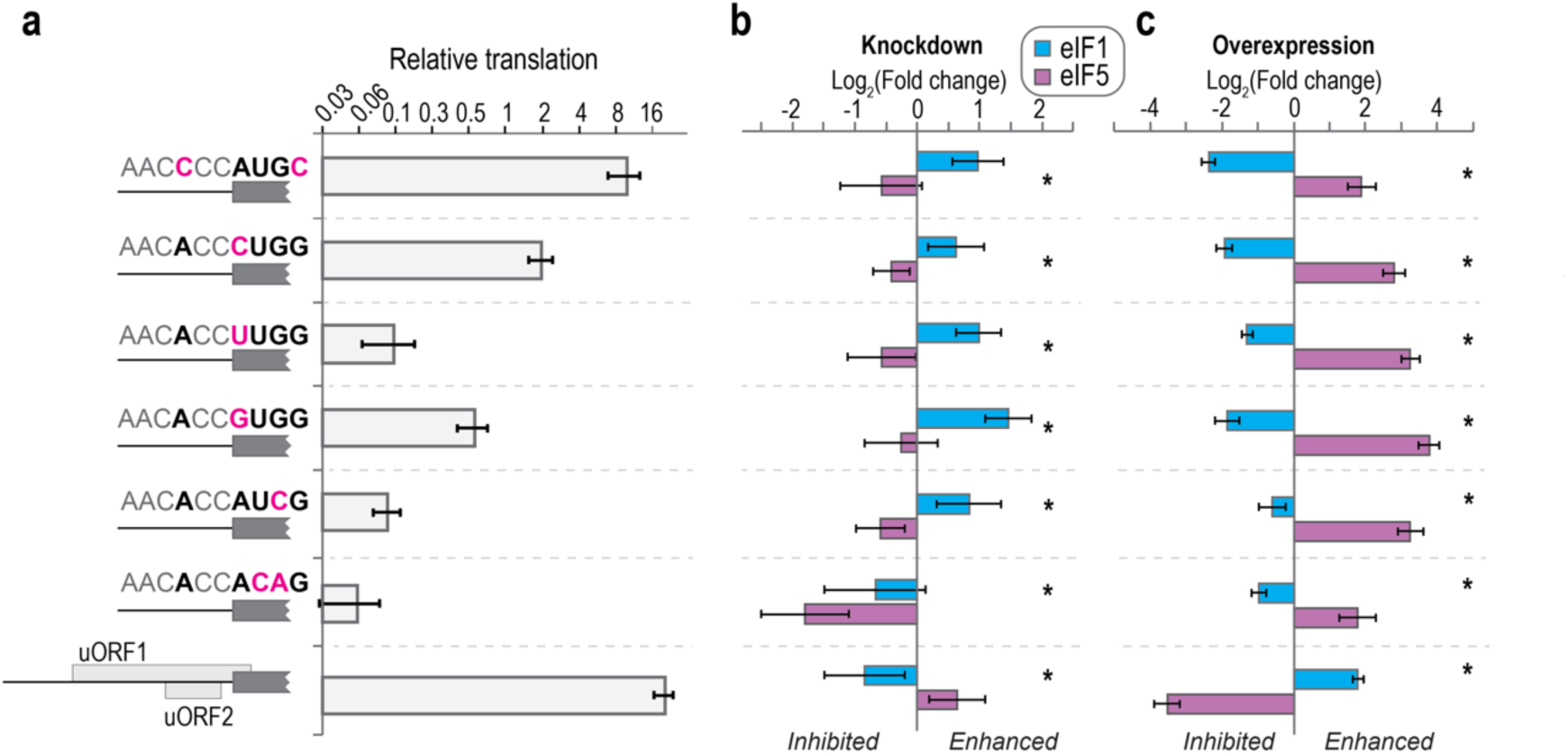
eIF1 and eIF5 have opposing roles in cells. (**a**) Plot of the relative translation activity of the indicated reporter mRNAs. The top 6 mRNAs contained unstructured 5’UTRs 50 nt in length, which were comprised of CAA repeates. For all 6, translation activity was quantified relative to model mRNA with an AUG in ideal Kozak context. The bottom mRNA was derived from the native 5’UTR of eIF5, which contains multiple upstream start sites in poor context. For this mRNA, translation activity was quantified relative to the same mRNA except the potential upstream start sites were eliminated. **(b)** Plot of the fold change of the translation activity of the indicated mRNAs upon siRNA-mediated knockdown of eIF1 and eIF5. **(c)** Plot of the fold change of the translation activity of the indicated mRNAs upon overexpression of eIF1 and eIF5. n = 5 and n = 6 for the knockdown and overexpression data, respectively. * indicates p < 0.01 for Student’s t-test (2 tailed) tests that compared eIF1 and eIF5 effects on the given mRNAs.

Alterations of eIF1 and eIF5 abundances differentially impacted translation of mRNAs that contain alternative start sites. As predicted, the presence of an alternative translation start site reduced translation of the reporter mRNA (**Fig. 5a**). However, relative to control cells, knockdown of eIF1 modestly enhanced translation of mRNAs with alternative start sites by about 2-fold, whereas eIF5 knockdown modestly inhibited translation of those same mRNAs (**Fig. 5b**). Strikingly, overexpression of either protein had the inverse effect. Overexpression of eIF1 inhibited translation of the mRNAs with alternative start sites by 2-4 fold, and eIF5 overexpression enhanced translation by 4-16 fold (**Fig. 5c**). In each case, eIF1 versus eIF5 effects were significantly different (t-test, p < 0.01). Translation of the eIF5 reporter responded inversely relative to the mRNAs with alternative start sites upon changing eIF1 and eIF5 abundance. This observation agrees with the known auto– and cross-regulation of eIF5 mRNA translation by eIF5 and eIF1, respectively, which is mediated by the inhibitory uORFs that have AUGs in poor context^37^.

## DISCUSSION

We defined a biophysical framework for ribosomal recognition of the translation start site during initiation (**Fig. 4d**). As it loads onto an mRNA, the 43S initiation complex contains the 40S ribosomal subunit, eIF1, eIF1A, and Met-tRNA_i_^Met^; each of these factors have been visualized directly as they load in real time (this study and ^22^). While currently invisible in our assays (a limitation of our study), extensive prior biochemical and structural data indicate that this complex also includes eIF2 and eIF3 (rev. in ^7–10,13,15^). By contrast, the 43S initiation complex lacked eIF5 as it loaded onto mRNA. Our finding explains the absence of eIF5 in structures of mammalian initiation complexes prior to recognition of the start site^44–53^. This divergence from yeast, where eIF5 incorporates into 43S initiation complexes^38,39^, may reflect the divergence of *S. cerevisiae* and human eIF3 (5 subunits versus 12). Regardless, such a divergence in kinetics has precedence in translation initiation. Universally-conserved eIF5B binds initiation complexes with inverted kinetics in yeast and humans, while still guiding joining of the 60S subunit in both organisms^19,22^.

eIF1 remained stably bound to the initiation complex until the start site was recognized. Our findings strongly suggest that this initial eIF1 binding event marks the scanning and initial start codon recognition steps. On ∼50 nt 5’UTRs, the initial eIF1 molecule remained bound for about two seconds on average. The duration increased to five seconds on the mRNA with the 200 nt unstructured 5’UTR. Conversely, eIF1 remained stably bound to initiation complexes on mRNAs without a translation start site. Initial recognition of the start site therefore very likely triggers initial eIF1 release, as was observed in studies of start site selection in yeast^23^. Consistently, the anticodon and stem of Met-tRNA_i_^Met^ shift by about 12 Å into the eIF1 binding site when paired properly with a start site (the P_scan_ to P_in_ transition) ^45,46,54–56^. The identical initial lifetimes of eIF1 on complexes with eIF2-GTP and eIF2-GDPNP indicate that this movement occurs independent of and likely prior to GTP hydrolysis by eIF2. Furthermore, we estimate a scanning rate of about 30 nt per s on the 200 nt model 5’UTR, after accounting for the eIF1 dissociation rate (3.1 ± 0.3 s^−1^) and assuming a 35 nt footprint of the initiation complex^61,62^. The increased population of initiation complexes with very stable initial eIF1 lifetimes on mRNAs with alternative start sites strongly suggests that many complexes bypass these non-ideal start sites.

After its initial dissociation upon recognition of the start site, eIF1 rapidly and transiently sampled initiation complexes. This period of rapid eIF1 reassociation likely provides additional opportunities to resume scanning at alternative or mis-identified translation start sites. Consistently, the sampling period and lifetimes of individual eIF1 binding events lengthened on mRNAs with alternative start sites. These effects were exaggerated on the mRNAs without a start site. Furthermore, experiments with unlabeled eIF1 (290 nM) reduced successful initiation on mRNAs with alterative start sites relative to experiments with labeled eIF1 (40 nM). We recapitulated our in vitro findings using both knockdown and overexpression studies in human cells. We therefore also conclude that changes in eIF1 abundance alters eIF1 occupancy on initiation complexes during the sampling window, which tunes the fidelity of start site recognition.

eIF5 only transiently binds initiation complexes. The very modest concentration dependence of eIF5 association indicates the presence of at least one rate-limiting upstream step that inhibits eIF5 association. Consistently, eIF5 reassociated at least 6-fold more rapidly on complexes with multiple eIF5 binding events. While currently impossible to elucidate in our assays, premature eIF5 association with initiation complexes may be prevented by both eIF1 occupancy and conformational rearrangements of eIF2 or subunits of eIF3 (e.g., eIF3c). Regardless, progression past the upstream inhibitory step(s) occurs independent of GTP hydrolysis, as eIF5 rapidly and continuously bound and released initiation complexes that contained eIF2-GDPNP. Furthermore, the 9 s delay before eIF5 productively bound at the highest concentration we examined (250 nM eIF5) matched the length of the eIF1 binding window (at 40 nM eIF1). The delay likely increased to about 14 s when unlabeled eIF1 and eIF5 were present at saturating, equimolar concentrations (290 nM) (see, **Figure S5**). Thus, eIF5 likely competes with eIF1 to bind initiation complexes near the end of the eIF1 sampling window. Only one or two eIF5 binding events are needed for the protein to trigger eIF2 dissociation and allow eIF5B association. The co-occupancy of eIF5 and eIF5B on initiation complexes suggests that a direct handoff of the Met-tRNA_i_^Met^ acceptor stem occurs from eIF2 to eIF5B.

eIF5 binding varied in response to the identity of the translation start site. eIF5 rarely bound initiation complexes on mRNAs with no start site or the AUC alternative site. eIF5 also bound initiation complexes on CUG, UUG, and GUG start sites much less frequently. On the subsets of initiation complexes that progressed to eIF5 and eIF5B association, we estimate that the apparent affinity of eIF5 is ∼2-fold lower for CUG start sites and at least 14-fold lower for UUG and GUG sites, which results from changes to both eIF5 association and dissociation rates. Our kinetic data agree with structures that identified direct contacts between eIF5 and the anti-codon stem and loop of Met-tRNA_i_^Met 55,56^. The destabilization of eIF5 on initiation complexes at alternative start sites inhibits progression to eIF5B association and ribosomal subunit joining. It also likely promotes eIF1 reassociation to resume scanning at non-ideal start sites. Thus, eIF5 binding represents a second bimolecular reaction during the otherwise unimolecular start codon recognition process that controls the fidelity of translation start site recognition.

Multiple kinetic steps – initial eIF1 release, eIF1 sampling, eIF5 binding, and eIF5 release – therefore underlie recognition of the translation start site. We propose that these kinetic steps couple to dynamic conformational rearrangements within the initiation complex (rev. in ^9,10,13,15^). During scanning, the initiation complex predominately populates a conformation with the anticodon of Met-tRNA_i_^Met^ disengaged from the mRNA (P_scan_). This state enables stable eIF1 binding. However, the complex also must engage (P_in_) and disengage (P_scan_) the anticodon with the mRNA on a microseconds to millisecond timescale to enable recognition of the start site during scanning. The presence of eIF1 and absence of base-pairing interactions between Met-tRNA_i_^Met^ and the mRNA rapidly shifts the complex back to the disengaged state (P_scan_). By contrast, proper recognition of the start site – and its stabilization from the newly formed three base-pair interactions between the initiator tRNA and start codon – lengthens the residence time of the complex in the engaged state (P_in_), which ejects the initial eIF1 protein. While presumably paused at the start site, the complex then rapidly samples multiple conformational states to engage and disengage Met-tRNA_i_^Met^ with the mRNA. These fluctuations permit stable reassociation of eIF1 to the disengaged state (P_scan_) and resumption of scanning on mis-identified start sites. During this period, eIF5 competes with eIF1 to capture the engaged conformation (P_in_) and allow initiation to proceed, which is enhanced on ideal start sites. The inverse effects by alternative start sites on eIF1 and eIF5 ribosome interactions (stabilized and destabilized, respectively) suggests that the energy barrier between the various conformational states is low; population of the states within the conformational landscape can be modulated by a single base pair. Our model also highlights that multiple offramps from the productive initiation pathway exist. Upon misidentification of a start site, complexes can either resume scanning (via stable eIF1 reassociation) or stall in states that allow eIF1 sampling but prevent eIF5 association.

Collectively, our biophysical model rationalizes more than a decade of genetic, biochemical, and structural data. The presence of multiple, competing bimolecular reactions on initiation complexes paused at potential start sites not only builds fidelity, but also allows flexible tuning of start site selection via changes in the levels of eIF1 or eIF5. Indeed, dynamic release of eIF1 from the nucleus is proposed to reprogram the cellular proteome during mitosis through differential usage of alternative start sites^63^. Our results underscore the dynamic plasticity of translation and its control in health and disease.

## Acknowledgements

We are grateful to members of the Lapointe, Dever, and Puglisi labs for helpful guidance, discussions, and feedback. We thank Peter Sarnow and the Sarnow lab (Stanford) for sharing cell culture equipment.

## Data, materials, and code availability

All single-molecule data are included in the manuscript. Raw files were processed with custom MATLAB code available on github (https://github.com/LapointeLab/eIF1-eIF5-2024-paper).

## METHODS

### Labeled eIF1

Synthetic DNA that encoded human eIF1 with C69 substituted for alanine was purchased from IDT and inserted into the 6His-MBP expression vector (v1C) from the UC Berkeley QB3 Macrolab. Using maleimide chemistry, this construct was used to conjugate Cy5 dye on the sole remaining cysteine94 residue in eIF1. The plasmid was transformed into Rosetta2 cells purchased from the UC Berkeley QB3 MacroLab and grown overnight at 37 °C on LB agar plates supplemented with 50 µg/mL kanamycin. Liquid cultures of single colonies were grown to OD_600_ β 0.5 at 37 °C in LB supplemented with kanamycin. After addition of 0.5 mM IPTG, the cultures were grown for 4 hours at 30 °C. Cells were harvested by centrifugation at 5,000 x *g* for 15 min at 4 °C in a Fiberlite F9 rotor (ThermoFisher, cat. # 13456093). Cells were lysed by sonication in lysis buffer (20 mM Tris-HCl pH 8.0, 300 mM NaCl, 10% (v/v) glycerol, 20 mM imidazole, and 5 mM β-mercaptoethanol), and lysates were cleared by centrifugation at 38,000 x *g* for 30 min at 4 °C in a Fiberlite F21 rotor followed by filtration through a 0.22 µm syringe filter. Clarified lysate was loaded onto a Ni-NTA gravity flow column equilibrated in lysis buffer, washed with 20 column volumes (CV) of lysis buffer, 20 CV of wash buffer (20 mM Tris-HCl pH 8.0, 1 M NaCl, 10% (v/v) glycerol, 40 mM imidazole, and 5 mM β-mercaptoethanol), and 10 CV of lysis buffer. Recombinant proteins were eluted with five sequential CV of elution buffer (20 mM Tris-HCl pH 8.0, 300 mM NaCl, 10% (v/v) glycerol, 300 mM imidazole, and 5 mM β-mercaptoethanol). Fractions with recombinant protein were identified by SDS-PAGE analysis. The relevant fractions were dialyzed overnight at 4 °C into TEV cleavage buffer (50 mM HEPES-KOH pH 7.5, 200 mM NaCl, 10% (v/v) glycerol, and 1 mM DTT) in the presence of excess TEV protease. TEV protease and the cleaved 6His-MBP tag were removed via a subtractive Ni-NTA gravity column equilibrated in TEV buffer, with the flow-through collected. The protein sample was diluted to 50 mM NaCl, applied to 1 mL SP HP ion exchange column, and eluted using a 50 mM to 500 mM NaCl gradient in the absence of reducing agents. Fractions that contained eIF1 at > 95% purity were concentrated to ∼ 22 µM, frozen in liquid N_2_, and stored at –80 °C. To fluorescently label the purified protein, we added 70-100 µL of purified protein (∼15 nmoles) to 1 mg of malemide-Cy5 in the presence of 2.5 mM TCEP. The reaction was incubated at room temperature for 20 min in the dark followed by an overnight incubation on ice. The unreacted Cy5 dye was removed using a 10DG-desalting column (Bio-Rad, cat.# 7322010) equilibrated in size-exclusion buffer (20 mM HEPES-KOH pH 7.5, 150 mM KOAc, 10% (v/v) glycerol, and 1 mM DTT). The labeled protein then was purified using a 23 mL SD75 size-exclusion chromatography column. Fractions that contained eIF1 at > 95% purity were concentrated to ∼ 10 µM (by Cy5), frozen in liquid N_2_, and stored at –80 °C.

### Labeled eIF5

Synthetic DNA that encoded human eIF5 with an N-terminal ybbR tag was purchased from IDT and inserted into the pET28 backbone, which contained a TEV protease cleavage site between the 6His-tag and the ybbR-tag. Rosetta2 cells were grown to OD 0.5 and induced with 0.5 mM IPTG for 4 hours at 30 °C. ybbR-eIF5 was purified essentially as above for eIF1, except 10 µM zinc sulfate was added to all buffers. Prior to ion exchange purification, ybbR-eIF5 was labeled with either Cy5 or Cy5.5 fluorescent dye using Sfp synthase as described. Briefly, ∼10 µM ybbR-eIF5 was supplemented with 10 mM MgCl_2_ and incubated at 37 °C for 90 min in the presence of 2-4 µM Sfp synthase enzyme and 20 µM of Cy5-CoA or Cy5.5-CoA substrate. Free dye was removed via purification over 10DG-desalting columns (Bio-Rad, cat.# 7322010) equilibrated in TEV cleavage buffer supplemented with 20 mM imidazole. TEV protease, Sfp synthase, and any remaining cleaved 6His tag were removed via a second subtractive Ni-NTA gravity column equilibrated in TEV buffer, with the flow-through collected. The labeled ybbR-eIF5 proteins were diluted to 50 mM NaCl and purified using a 1 mL Q HP column. eIF1 was eluted using a 50 mM to 500 mM gradient of NaCl. Fractions that contained ybbR-eIF5 at > 95% purity were concentrated to ∼ 10 µM (by dye; ∼50% labeled), frozen in liquid N_2_, and stored at –80 °C.

### Labeled eIF5B

N-terminally truncated (587-1220) eIF5B with an N-terminal ybbR-tag was purified and labeled with Cy3.5 dye using Sfp synthase as described^22^.

### Labeled initiator tRNA

HPLC purified, synthetic human Met-tRNA_i_ with U46 replaced with Uridine-C6 Amino linker conjugated to Cyanine 3 SE was purchased from TriLink. Prior to aminoacylation, the fluorescently-labeled tRNA_i_-Cy3 was purified using phenol:chloroform extraction and ethanol precipitation. Aminoacylation with methionine was performed as described^41^. The charging efficiency was > 90% based on acid urea PAGE analyses.

### Unlabeled eIFs

Recombinant eIF1^64^, eIF1A^64^, eIF3j^64^, eIF4AI^64,65^, eIF4B^22^, eIF4G_165-1599_^22^, eIF4E^65,66^, eIF5^41^ proteins were purified as described. Native human eIF2 and eIF3 were purified as described^64,67^. Importantly, purified human eIF3 lacked any detectable presence of eIF1 or eIF5 by mass spectrometry analysis (**Table S2**). Unlabeled human Met-tRNA_i_^Met^ was in vitro transcribed and aminoacylated as described^22^.

### Ribosomal subunits

Human 40S and 60S ribosomal subunits were purified from either wild-type or the appropriately edited HEK293T cell lines (RPS15-ybbR for 40S-Cy3) and (RPL5-ybbR for 60S Cy5/Cy5.5) and labeled with fluorescent dyes as described^57,68^. Cell line identities were confirmed by PCR assays, Sanger sequencing, western blotting, and fluorescence gels (after labeling). Cells were not tested for mycoplasma contamination.

### mRNAs

Full-length human β-globin mRNA (NM_000518.5, 628 nts) was in vitro transcribed and prepared as described^22^ to yield a 5’ m^7^G capped and 3’-biotinylated mRNA with a 30 nt poly(A) tail. To generate the short, unstructured model RNAs, DNA that encoded the reverse complement of the RNA and T7 promoter sequences were purchased from IDT as single-stranded oligos. After hybridizing a second oligo that encoded the sense T7 promoter sequence, the RNAs were in vitro transcribed using T7 RNA polymerase for 3 hrs at 37 °C and treated with Turbo DNase. The transcribed RNAs were prepared identically to β-globin mRNA, except the RNAs lacked the poly(A) tail and all purifications were done using standard phenol:chloroform extractions and ethanol precipitations. To generate the mRNA with a 200 nt unstructured 5’UTR, we replaced the 5’UTR on β –globin mRNA with 198 nt of CAA repeats, leaving only the Kozak context around the AUG (CACCAUGGA). The mRNA was in-vitro transcribed and prepared identical to β –globin, producing a 5’-m7G capped, 3’-biotinylated mRNA with a 202 nt 5’UTR and a 30 nt poly(A) tail.

### Real-time single-molecule analyses

All real-time imaging was conducted using a modified Pacific Biosciences RSII DNA sequencer and the Maggie software (v. 2.3.0.3.154799), which were described previously^58^. All experiments were performed at 30 °C (unless otherwise noted) using a 532 nm excitation laser at 0.32 µW/µm^2^, which directly excited Cy3 and Cy3.5 dyes. Cy5 and Cy5.5 dyes were excited either directly via a 642 nm laser (0.1 µW/µm^2^) or via FRET as indicated. Four-color fluorescence emission (Cy3, Cy3.5, Cy5, and Cy5.5) was detected at 10 frames per second for 600 s. ZMW chips were purchased from Pacific Biosciences, which were passivated by reaction with polyvinylphosphonic acid to form a covalent Al-polyphosphonate coating 60. Prior to imaging, all ZMW chips were washed with 0.2% Tween-20 and TP50 buffer (50 mM Tris-OAc pH 7.5, 100 mM KCl). Washed chips were coated with neutravidin by a 5 min incubation with 1 µM neutravidin diluted in TP50 buffer supplemented with 0.7 mg mL^−1^ UltraPure BSA and 1.3 µM of pre-annealed DNA blocking oligos (CGTTTACACGTGGGGTCCCAAGCACGCGGCTACTAGATCACGGCTCAGCT) and (AGCTGAGCCGTGATCTAGTAGCCGCGTGCTTGGGACCCCACGTGTAAACG). The imaging surface then was washed with TP50 buffer at least four times. All experiments, which are outlined below, were performed essentially as described^22^.

The ‘initiation reaction buffer’ was: 20 mM HEPES-KOH, pH 7.5, 70 mM KOAc, 2.5 mM Mg(OAc)_2_, 0.25 mM spermidine, 0.2 mg mL^−1^ creatine phosphokinase, 1 mM ATP•Mg(OAc)_2_, and 1 mM GTP•Mg(OAc)_2_. The ‘imaging buffer’ was the initiation reaction buffer supplemented with casein (62.5 µg mL-1), 5 mM TSY, and an oxygen scavenging system^69^: 2 mM protocatechuic acid (PCA) and 0.06 U/µL protocatechuate-3,4-dioxygenase (PCD).

To prepare the eIF2–GTP–Met-tRNA_i_^Met^ ternary complex (TC), 3.3 µM eIF2 was incubated in initiation reaction buffer (excluding ATP•Mg(OAc)_2_) for 10 min at 37 °C to saturate eIF2 with GTP. The eIF2– GTP complex then was incubated with either 2.3 µM of unlabeled Met-tRNA_i_^Met^ (as a control) or Met-tRNA_i_^Met^-Cy3 (tRNA_i_-Cy3) for 5 min at 37 °C to form TC. In a few instances, GTP was replaced with a non-hydrolyzable analog (GDPNP) during formation of TC, which prevents GTP hydrolysis and subsequent departure of eIF2 from the initiation complex.

To prepare the 43S pre-initiation complex (43S PIC), we typically incubated 1 µM eIF1, 1 µM eIF1A, 500 nM TC (by eIF2), 1 µM eIF5, 400 nM eIF3, 1.2 µM eIF3j, and 240 nM 40S subunits for 5 min at 37°C in initiation reaction buffer. In most experiments with eIF1-Cy5 or eIF5-Cy5/5.5, the unlabeled protein was replaced with the labeled version at the same concentration (1 µM, determined via dye fluorescence). The exception was for the eIF5-Cy5.5 titration experiments (**Fig. 3**); for this, eIF5-Cy5.5 was included at 62 nM, which was increased to the reported 10, 20, 40, 80, and 250 nM final concentration by inclusion of additional eIF5-Cy5.5 in the 43S PIC mixture below. Finally, in a few experiments (**Fig. 1**), the eIF5 concentration was reduced to 125 nM to yield the final concentration of 5 nM.

In the real-time single-molecule initiation assays, the indicated 5’ m^7^G capped and 3’-biotinylated mRNA was tethered to a prepared, neutravidin-coated ZMW imaging surface. Immediately after tethering and washing the surface with imaging buffer, a mixture of 2 µM eIF4A, 440 nM eIF4B, 260 nM eIF4G, and 320 nM eIF4E in 20 µL of imaging buffer was added to the surface. The surface and reaction mixture were incubated at room temperature for 5-10 minutes as the instrument initialized. At the start of data acquisition, a 20 µL mixture of 10 nM 43S PIC (by the 40S subunit), 80 nM eIF5B-Cy3.5, and 300 nM 60S subunits in imaging buffer supplemented with 500 nM of unlabeled eIF1, eIF1A, and/or eIF5 was added to the surface. For the eIF5-Cy5.5 titration experiments, additional eIF5-Cy5.5 was included in this mix to obtain the reported final concentrations. Importantly, addition of this second reaction mixture to the surface doubled the total reaction volume (to 40 µL; since 20 µL with eIF4 proteins was already present); thus, unless indicated otherwise, the final concentrations of individual components in the experiments were: 1 µM eIF4A, 220 nM eIF4B, 230 nM eIF4G, 320 nM eIF4E, 10 nM 40S subunits, 20 nM TC, 16 nM eIF3, 48 nM eIF3j, 290 nM eIF1A, 290 nM eIF1 (unlabeled), and 290 nM eIF5 (unlabeled). In experiments with labeled eIF1-Cy5 and eIF5-Cy5.5, the unlabeled versions of the proteins were excluded from all steps and solutions.

### Equilibrium TIRF microscopy

A home-built, prism-based TIRFM instrument has been described previously^69–71^. Emission data were collected in both the Cy3 (donor) and Cy5 (acceptor) channels following excitation of the Cy3 donor dye with the 532 nm laser. All complexes were prepared as above, and TC-GDPNP-tRNA_i_-Cy3 was used. The pre-assembled 43S PIC (75 nM) was incubated with the indicated biotinylated model mRNA (AUG or No AUG at 50 nM) for 15 min at 37 °C in the initiation reaction buffer. The mRNA-43S complex was tethered to the neutravidin coated imaging surface for 5 min at room temperature at 5 nM final concentration (by 40S subunits). Following a 50 µL wash of the surface using initiation reaction buffer, the sample was incubated with the imaging buffer supplemented with 300 nM eIF1A and either 20 nM eIF1-Cy5 or 20 nM eIF5-Cy5, as indicated. Emission data were collected in both the Cy3 (donor) and Cy5 (acceptor) channels following excitation of the Cy3 donor dye with the 532 nm laser. Fluorescence intensities from single complexes were analyzed as outlined below.

### Data analyses

Experimental movies that captured fluorescence intensities over time were processed using MATLAB R2018-b as described previously. In all analyses, spots (equilibrium TIRF) or ZMWs (real-time analyses) with the desired fluorescence signals were identified by filtering for the desired signals.

To determine *E_FRET_*, fluorescence data from individual ZMWs with the desired fluorescence signals were background corrected using SPARTAN 3.7.0^72^, with single molecules indicated by single-step photobleach events of the donor fluorophore. FRET on and off states were assigned automatically using vbFRET (version june10)^73^, which were visually inspected and manually corrected as needed. An *E_FRET_* threshold of 0.1 was used in all experiments. *E_FRET_* was defined as:

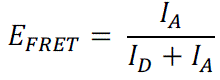

Where *I_D_* and *I_A_* represent fluorescence intensities of the donor and acceptor fluorophores. *E_FRET_* values observed across all events and molecules were binned (50 bins, –0.2 to 1.2) and fit to gaussian functions to determine the mean and standard deviations.

To determine association and dissociation kinetics, binding events of individual components were assigned manually based on the appearance and disappearance of the respective fluorescence signals. The observed times for an event to occur from 100-200 individual molecules (i.e., ZMWs) were used to calculate cumulative probability functions of the observed data (cdfcalc, MATLAB), which were fit to single-or double-exponential functions in MATLAB (cftool, non-linear least squares method) as described. All derived association rates, median association times, lifetimes, and the number of molecules examined are reported in **Tables S1 and S3**. The exponential function was defined as:

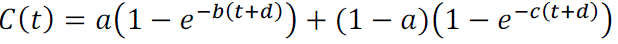

where *t* is time (in s), *b* and *c* rates, and *d* an adjustment factor. If a parameter yielded a phase that represented less than 10% of the population, only a single phase was used to derive the respective rate and reported (i.e., single-exponential function). For eIF5, we focused kinetic analyses on the fastest 90% of observed molecules to enable fits to single-exponential functions and facilitate comparisons.

To facilitate visualization of events in main text figures, raw fluorescent intensities were manually corrected to account for spectral bleedthrough across the various emission channels. Reported errors for derived rates represent 95% C.I. yielded from fits to linear, single-exponential, or double-exponential functions, as indicated.

### eIF3 sample preparation for mass spectrometry analysis

The eIF3 protein solutions were brought up to 50 μL with 40 μL 50 mM ammonium bicarbonate. The solutions were reduced with dithiothreitol (10 mM final in 50 mM ammonium bicarbonate) at 56 °C for 45 min. The solutions were alkylated with 2-chloroacetamide (20 mM final in 50 mM ammonium bicarbonate) and incubated in the dark at ambient temperature for 30 min. Trypsin (Promega, Madison, WI) 250 ng, was added and the solutions were incubated overnight at 37 °C with mixing. Samples were allowed to equilibrate to ambient temperature and were taken to dryness on a speedvac. The dried samples were brought up in 30 μL 85% Acetonitrile/15 mM ammonium formate pH 2.8. The peptide solution was desalted over 1cm TopTip packing material (Poly-LC-, Columbia, MD) packed in a 10 μL pipette tip. Material was washed with 50 μL 85% Acetonitrile/15 mM ammonium formate pH 2.8, 3 times and eluted with 50 μL 15 mM ammonium formate pH 2.8. The desalted material was taken to dryness in a speed vac.

### Orbitrap Fusion LC/MS/MS analysis

Desalted samples were brought up in 2% acetonitrile in 0.1% formic acid (20 μL) and were analyzed (2 μL) by LC/ESI MS/MS with a Thermo Scientific Easy-nLC 1200 (Thermo Scientific, Waltham, MA) nano HPLC system coupled to a tribrid Orbitrap Fusion (Thermo Scientific, Waltham, MA) mass spectrometer. In-line de-salting was accomplished using a reversed-phase trap column (100 μm × 20 mm) packed with Magic C18AQ (5-μm 200Å resin; Michrom Bioresources, Bruker, Billerica, MA) followed by peptide separations on a reversed-phase column (75 μm × 270 mm) packed with Reprosil C18AQ (3-μm 100 Å resin; Dr. Maisch, Germany) directly mounted on the electrospray ion source and warmed to 37 °C with a column heater. A 45-minute gradient from 9% to 44% B at a flow rate of 300 nL/minute was used for chromatographic separations, with A as 0.1% formic acid and B 80% acetonitrile in 0.1% formic acid. The heated capillary temperature was set to 300 °C and a static spray voltage of 2200 V was applied to the electrospray tip. The Orbitrap Fusion was operated in the data-dependent mode, switching automatically between MS survey scans in the Orbitrap (AGC target value 500,000, resolution 120,000, and maximum injection time 50 milliseconds) with MS/MS spectra acquisition in the linear ion trap using quadrupole isolation. A 3 second cycle time was selected between master full scans in the Fourier-transform (FT) and the ions selected for fragmentation in the HCD cell by higher-energy collisional dissociation with a normalized collision energy of 27%. Selected ions were dynamically excluded for 45 seconds and exclusion mass by mass width +/-10 ppm.

### LC/MS/MS Data Analysis

Data analysis was performed using Proteome Discoverer 3.1 (Thermo Scientific, San Jose, CA). The data were searched against Uniprot Human (Uniprot UP000005640 Nov 11, 2023) (http://www.thegpm.org/crap/) fasta files. Trypsin was set as the enzyme with maximum missed cleavages set to 2. The precursor ion tolerance was set to 10 ppm and the fragment ion tolerance was set to 0.6 Da. Variable modifications included oxidation on methionine (+15.995 Da). Dynamic modifications on the protein N-terminus included acetylation (+42.011 Da), methylation (+14.016 Da) and methionine loss plus acetylation (–89.030 Da). Static modifications included carbamidomethyl on cysteine (+57.021 Da). Data were searched using Sequest HT. All search results were run through Percolator for scoring and identified peptides were filtered for 1% peptide-level false discovery rate using q value of 0.01.

### Cell Culture and Transfections

U2OS cells were kindly provided by Nancy Kedersha (Harvard University). U2OS cells were maintained in DMEM (Corning) supplemented with 1 mM L-glutamine, 1 mM sodium pyruvate, 10% FBS (Gibco) and penicillin/streptomycin (Quality Biological) at 37°C in 5% CO_2_. For luciferase assays, U2OS cells were grown overnight in 10 cm plates to ∼70% confluence, washed, treated with trypsin, and then transfected using Lipofectamine 2000 reagent (Invitrogen) in a one-day protocol in which suspension cells were added directly to the DNA mixtures in 96-well half-area white plates (Costar). For eIF1/eIF5 overexpression experiments, 0.2 µl Lipofectamine, 10 ng *Renilla* control reporter, and 10 ng firefly test reporter plasmid were mixed with 25 ng eIF1 or eIF5 overexpressing plasmid or empty control plasmid in 25 µl Opti-MEM (Gibco) and then dispensed to each well along with 10^4^ cells suspended in 25 µl DMEM. The transfection was terminated 24 h later by removing the media and lysing the cells using 25 µl 1x Passive Lysis Buffer (Promega). For eIF1/eIF5 knockdown experiments, 0.2 µl Lipofectamine, 5 ng *Renilla* control reporter, and 5 ng firefly test reporter plasmid were mixed with 25 ng mixture of eIF1 (four different plasmids, two each targeting EIF1 and EIF1B respectively, with every shRNA plasmid contributing 6.25 ng DNA to the total), eIF5 (two different plasmids, with every shRNA plasmid contributing 12.5 ng DNA to the total) or control (shc002) shRNA expressing plasmids in 25 µl Opti-MEM (Gibco) and then dispensed to each well along with 10^4^ cells suspended in 25 µl DMEM. The transfection was terminated 48 h later by removing the media and lysing the cells using 25 µl 1x Passive Lysis Buffer.

### Plasmids

Leader sequences matching those of the single-molecule *in vitro* experiments were synthesized and cloned between the *Hin*dIII and *Bam*HI restriction sites of the vector p2luc^74^ by LifeSct LLC (Rockville, MD). The sequence of the WT leader insert is: 5’-*`aagctt`*GGACAACAACAACAAACCCTCGAACAACAACAACAACAACAACAAC**A**CC**ATGG**ACAACAcCAA

CAcCAcC*ggatcc*-3’ (*Hin*dIII and *Bam*HI sites in italics, nucleotides in bold were mutated as shown in **Fig. 5a** to generate alternative start sites). The *Renilla* expressing plasmid used in these experiments, pSV40-*Renilla*, as well as the phRL-based eIF1 and eIF5 overexpressing plasmids and empty vector control plasmid were described previously^37^. The shRNAs for eIF1 and eIF5 knockdown experiments were purchased from Sigma. Each mRNA was targeted by two shRNAs – TRCN0000310709 and TRCN0000303471 targeting eIF1, TRCN0000149959 and TRCN0000149658 targeting eIF1B, and TRCN0000338465 and TRCN0000338404 targeting eIF5. For control, shc002 shRNA was used. The firefly reporters with the eIF5 wild-type and deregulated control leaders were described previously^37^.

### Dual Luciferase Assay

Luciferase activities were determined by the Dual Luciferase Stop and Glo Reporter assay system (Promega). Relative light units were measured on a CentroXS3 LB960 microplate luminometer fitted with two injectors (Berthold Technologies). Light emission was measured after injection of 25 µL firefly luciferase substrate followed by the same volume of *Renilla* luciferase substrate. The firefly luciferase activity was normalized relative to the activity of a plasmid expressing *Renilla* luciferase from an AUG codon in perfect context^37^. All reported p-values are derived from Student’s t-tests (2 tailed, type III).

---

**Figure S1.**
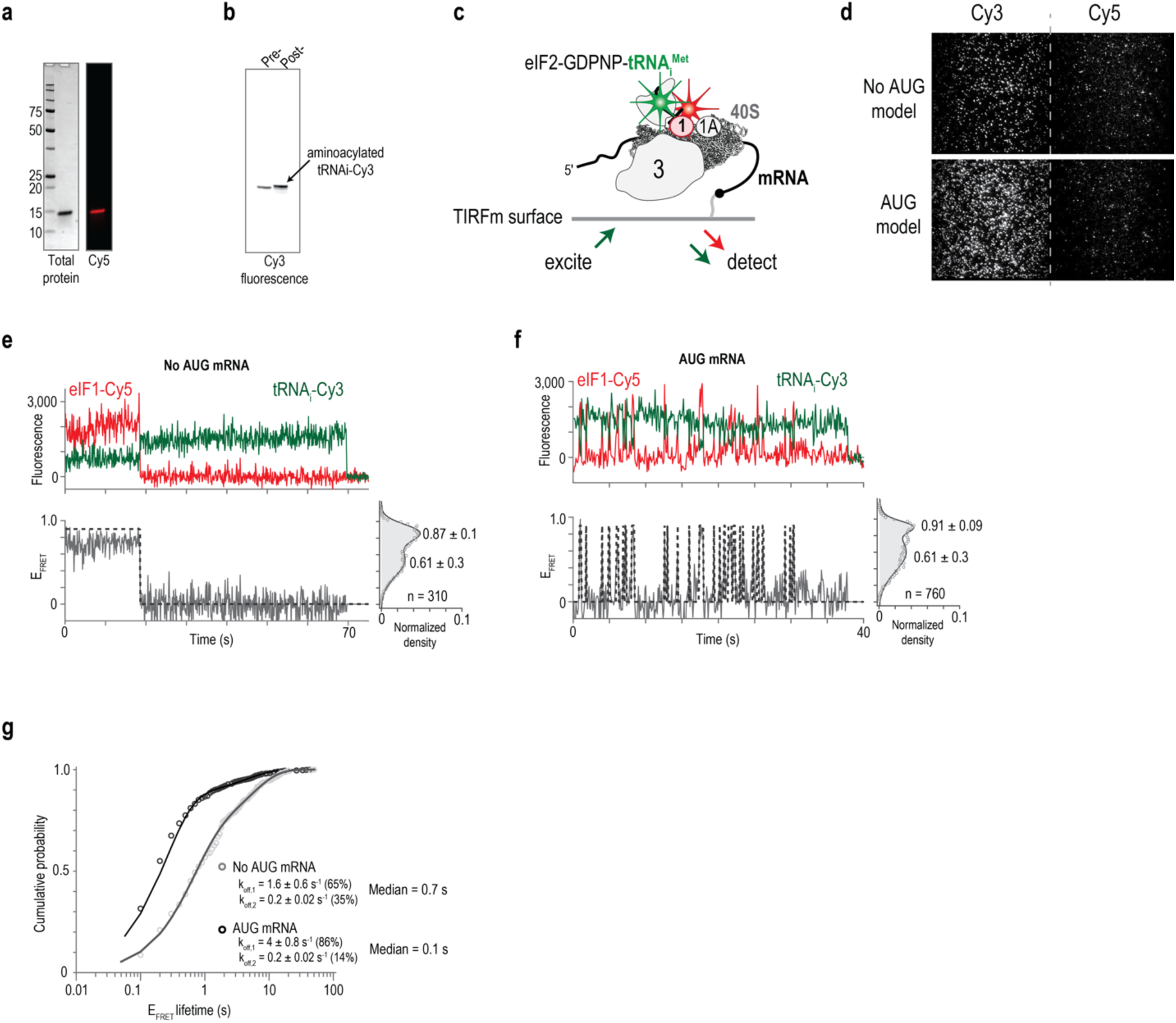
An eIF1 FRET signal. (**a**) Image of a gel from SDS-PAGE analysis of purified eIF1 labeled on C94 with Cy5 fluorescent dye. The left and right images are total protein and Cy5 fluorescent scans of the gel. **(b)** Image of an acid TBE-Urea gel that examined Met-tRNA_i_-Cy3 before and after in vitro aminoacylation. The gel was scanned for Cy3 fluorescence. **(c)** Schematic of the equilibrium total internal reflection fluorescence microscopy (TIRFm) experiments. Samples were excited with a 532 nM laser. **(d)** Example fields of view of TIRFm experiments. **(e,f)** Example fluorescence data of initiation complexes equilibrated on the model mRNA without a start site (No AUG model, panel e) or with a start start site (AUG model, panel f). The complexes contain tRNA_i_-Cy3 (green) to eIF1-Cy5 (red) FRET events. The top plots represent the fluorescence intensities (arbitrary units) of both signals, and the bottom plots represent the calculated FRET efficiency (E_FRET_). The right panels plot the E_FRET_ distribution observed on the indicated number of eIF1 binding events. **(g)** Cumulative probability plot of the eIF1 FRET lifetime observed on the two model mRNAs. The lines represent fits to double-exponential functions, which were used to derive the indicated rates.

**Figure S2.**
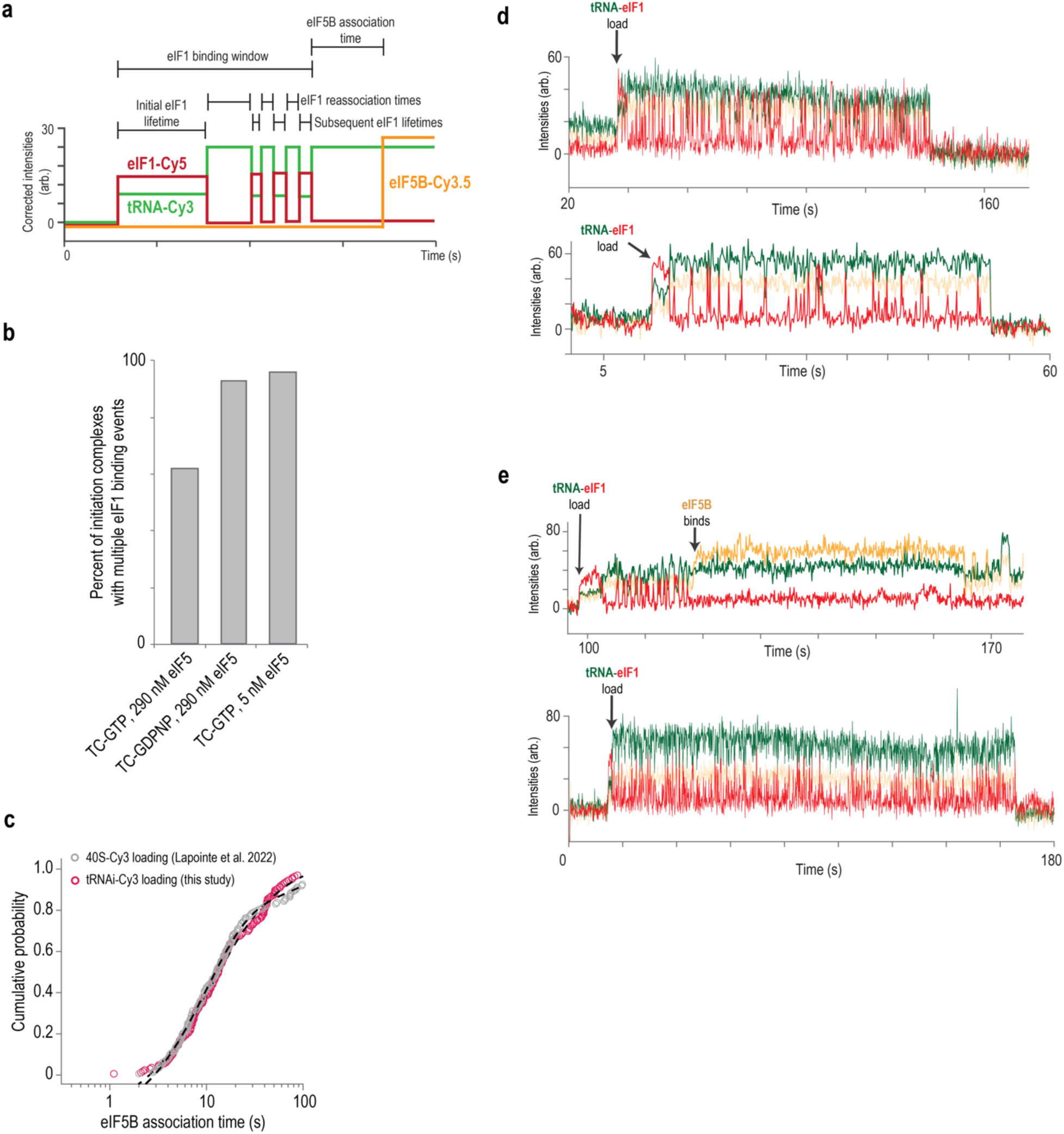
eIF1 stably binds and dynamically samples individual complexes. (**a**) Schematic of predicted single-molecule data for eIF1 experiments. Initial appearance of tRNA_i_-Cy3 (green) to eIF1-Cy5 (red) FRET signifies that an initiation complex loaded onto the tethered mRNA. This initial FRET event is defined as the ‘initial eIF1 binding event’ and the duration of this event defined as the ‘initial eIF1 lifetime’. The complex then contains multiple subsequent tRNA_i_-Cy3 to eIF1-Cy5 FRET events, which are defined as subsequent eIF1 binding events. The dwell time between loss of the previous eIF1 signal and appearance of the next eIF1 signal is defined as the ‘eIF1 reassociation time’. The collective duration of all subsequent eIF1 binding events is defined as the ‘subsequent eIF1 lifetime’. Appearance of eIF5B-Cy3.5 (orange) signal signifies successful entry to the final initiation steps that culminate with joining of the 60S ribosomal subunit. The dwell time from loss of the final eIF1-Cy5 signal to appearance of eIF5B-Cy3.5 signal is defined as the ‘eIF5B association time’. **(b)** Plot of the percentage of loaded initiation complexes that contain multiple eIF1 binding events on β-globin mRNA in the indicated conditions. **(c)** Cumulative probability plot of the eIF5B association time measured relative to initial loading of the initiation complex, which was signified by either appearance of tRNA_i_-Cy3 to eIF1-Cy5 FRET (this study) or initial appearance of 40S-Cy3 signal. The overlapped plots indicate that labeled tRNA_i_-Cy3 and eIF1-Cy5 function analogously as their unlabeled versions. **(d)** Example single-molecule data of eIF1 continuously sampling initiation complexes that contain eIF2-GDPNP, which are unable to progress to eIF5B association. **(e)** Example single-molecule data of eIF1 extensively sampling initiation complexes in the presence of a limiting concentration of eIF5 (5 nM). One complex slowly progressed to eIF5B association (top) and the other had continuous eIF1 sampling without eIF5B association.

**Figure S3.**
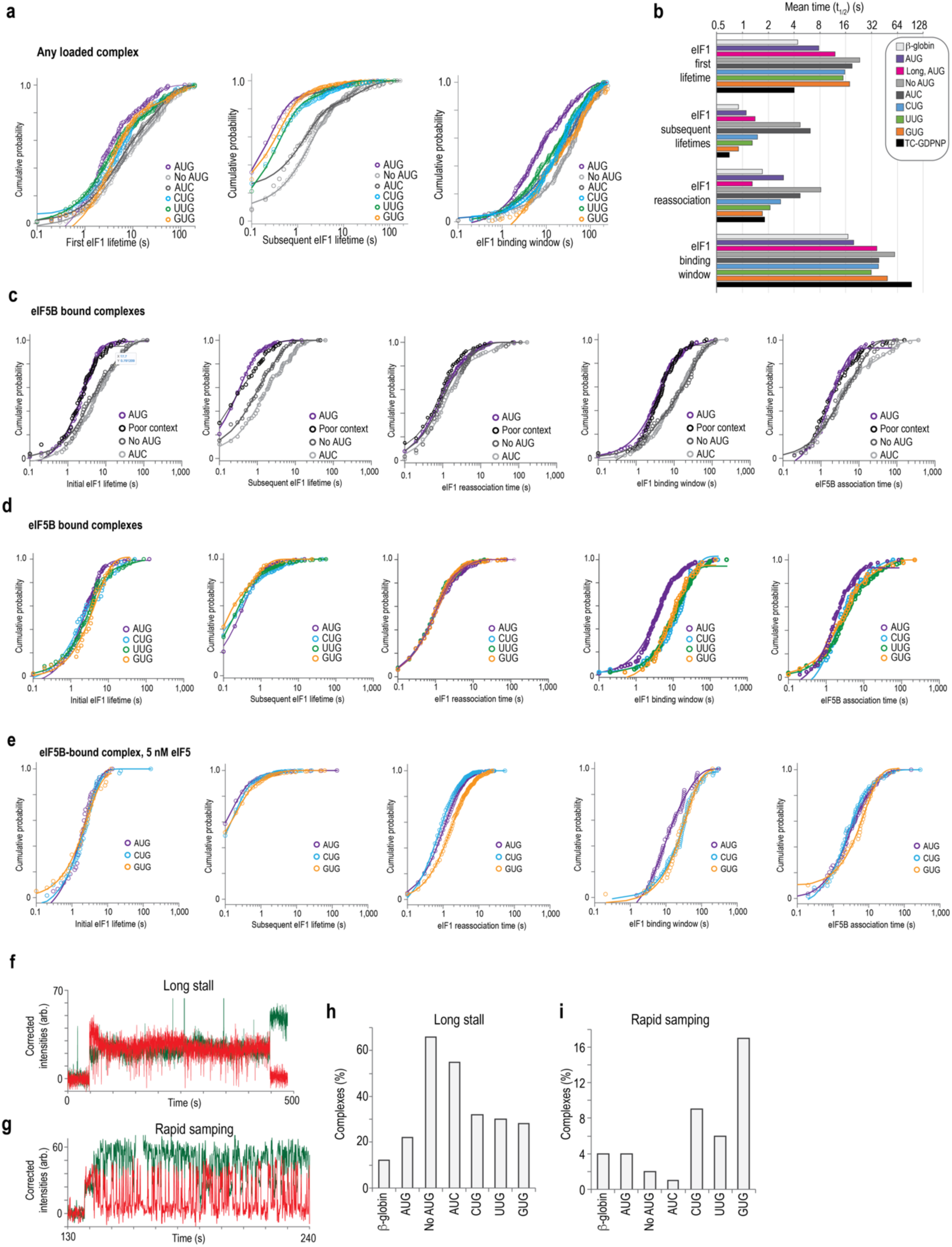
eIF1 kinetics depend on 5’UTR length and the identity of the translation start site. (**a**) Cumulative probability plots of the indicated eIF1 parameters observed on any loaded initiation complexes (both unsuccessful and successful). Lines represent fits to exponential functions. **(b)** Plot of the mean elapsed times (t_1/2_ values) for the indicated eIF1 parameters on the indicated mRNAs. All values plotted here were derived from events that occurred on any loaded initiation complex (both unsuccessful and successful). **(c)** Plot of the indicated kinetic parameters on eIF5B-bound initiation complexes (successful) on the model AUG mRNAs in ideal (**A**CC**<u>AUG</u>G**A) or poor (**C**CC**<u>AUG</u>C**A) Kozak context. **(d,e)** Cumulative probability plots of the indicated kinetic parameters on eIF5B-bound initiation complexes on the indicated model mRNAs in the presence of 290 nM (panel d) or 5 nM (panel e) unlabeled eIF5. Lines represent fits to exponential functions. **(f,g)** Example single-molecule data where the initiation complex stalled in either a state that permitted stable eIF1 binding (panel f) or allowed rapid eIF1 sampling (panel g). **(h,i)** Plots of the percent of all loaded initiation complexes that stalled in either the stable eIF1 binding state (panel h) or the rapid sampling state (panel i). In all experiments, eIF1-Cy5 was present at a final concentration of 40 nM (by Cy5) and unlabeled eIF5 was present at either 290 or, when indicated, 5 nM. See **Table S1** for the number of complexes and binding events analyzed in each experiment.

**Supplementary Figure S4.**
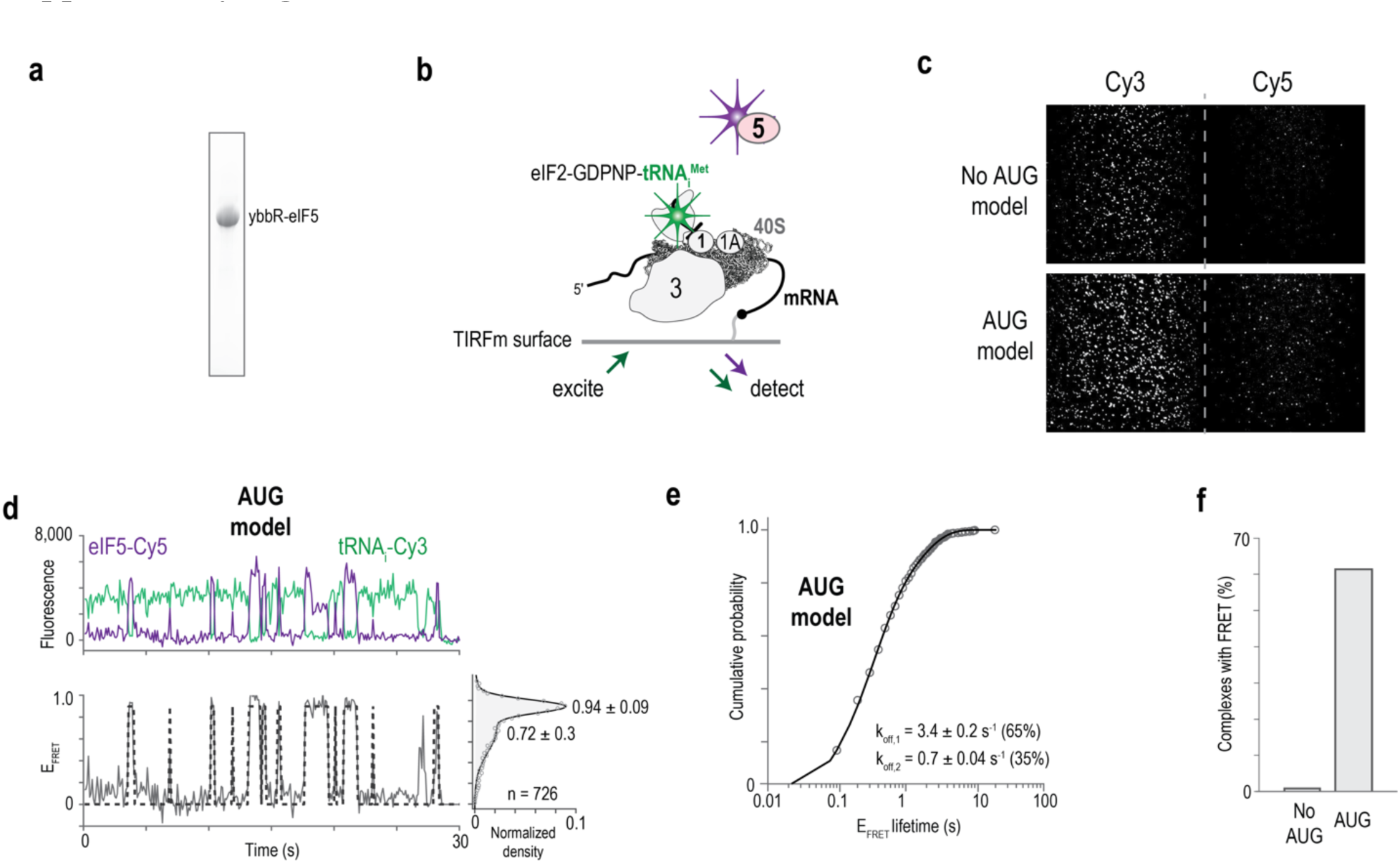
An eIF5 FRET signal. (**a**) Image of a gel from SDS-PAGE analysis of purified eIF5 tagged on its N-terminus with the ybbR peptide. **(b)** Schematic of the equilibrium total internal reflection fluorescence microscopy (TIRFm) experiments. Samples were excited with a 532 nM laser. **(c)** Example fields of view of TIRFm experiments. **(d)** Example fluorescence data of initiation complexes equilibrated on the model mRNA with a start start site. The complexes contain tRNA_i_-Cy3 (green) to eIF5-Cy5 (purple) FRET events. The top plot represents the fluorescence intensities (arbitrary units) of both signals, and the bottom plot represents the calculated FRET efficiency (E_FRET_). The right panel plots the E_FRET_ distribution observed on the indicated number of eIF5 binding events. **(e)** Cumulative probability plot of the eIF5 FRET lifetime observed on the AUG model mRNA. The line represent a fit to a double-exponential functions, which was used to derive the indicated rates. **(f)** Plot of the percent of initiation complexes that contained an eIF5 binding event on the indicated model mRNAs.

**Supplementary Figure S5.**
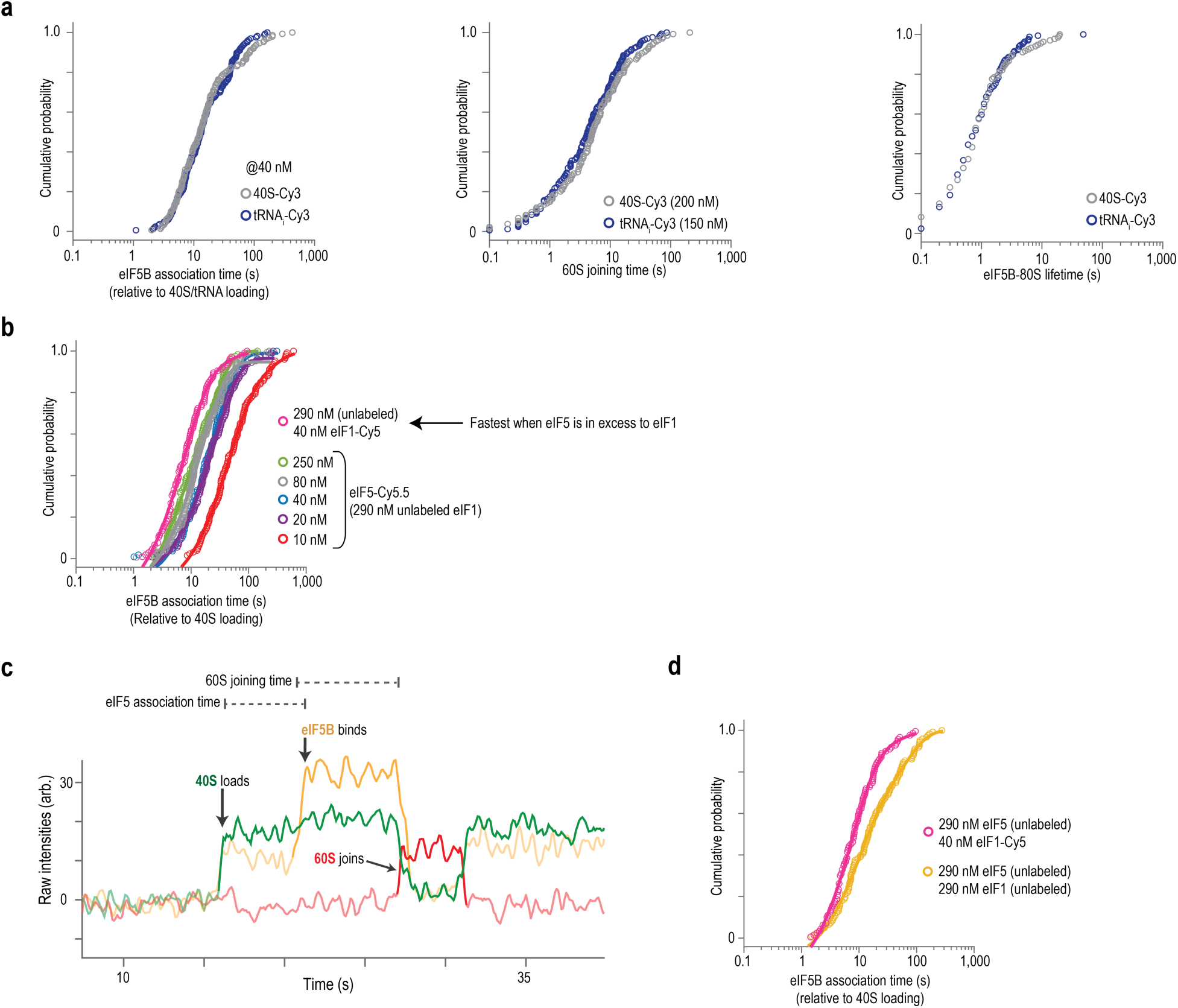
eIF5-Cy5.5 functions analogously to its unlabeled version. (**a**) Cumulative probability plot of the eIF5B association time, 60S subunit joining time, and the lifetime of eIF5B on the 80S initiation complex. Here, the eIF5B association time was measured relative to initial loading of the initiation complex, which was signified by either appearance of tRNA_i_-Cy3 signal (this study) or initial appearance of 40S-Cy3 signal^22^. The overlapped plots indicate that labeled tRNA_i_-Cy3 and eIF5-Cy5.5 function analogously as their unlabeled versions. **(b)** Cumulative probability plot of the eIF5B association times measured relative to loading of the 43S initiation complex, as signaled by appearance of tRNA_i_-Cy3 fluorescence. The magenta data (290 nM unlabeled eIF5, 40 nM eIF1-Cy5) is from the labeled eIF1 experiments. The remainder are from the labeled eIF5 experiments at the indicated concentrations. **(c)** Example single-molecule fluorescence data from an experiments that monitored loading of the 43S complex as signaled by 40S-Cy3 (green) in the presence of 40 nM eIF5B-Cy3.5 (orange), and 60S-Cy5 (red). In this experiment, unlabeled eIF1 and eIF5 were present at 290 nM each. **(d)** Cumulative probability plot of the eIF5B association times measured relative to loading of the 43S initiation complex, as signaled by appearance of tRNA_i_-Cy3 fluorescence (pink data, same data as in panel b) or 40S-Cy3 fluorescence (orange data). The concentration of eIF1 and eIF5 in the two experiments are indicated. See **Table S3** for the number of complexes and binding events analyzed in each experiment.

**Supplementary Figure S6.**
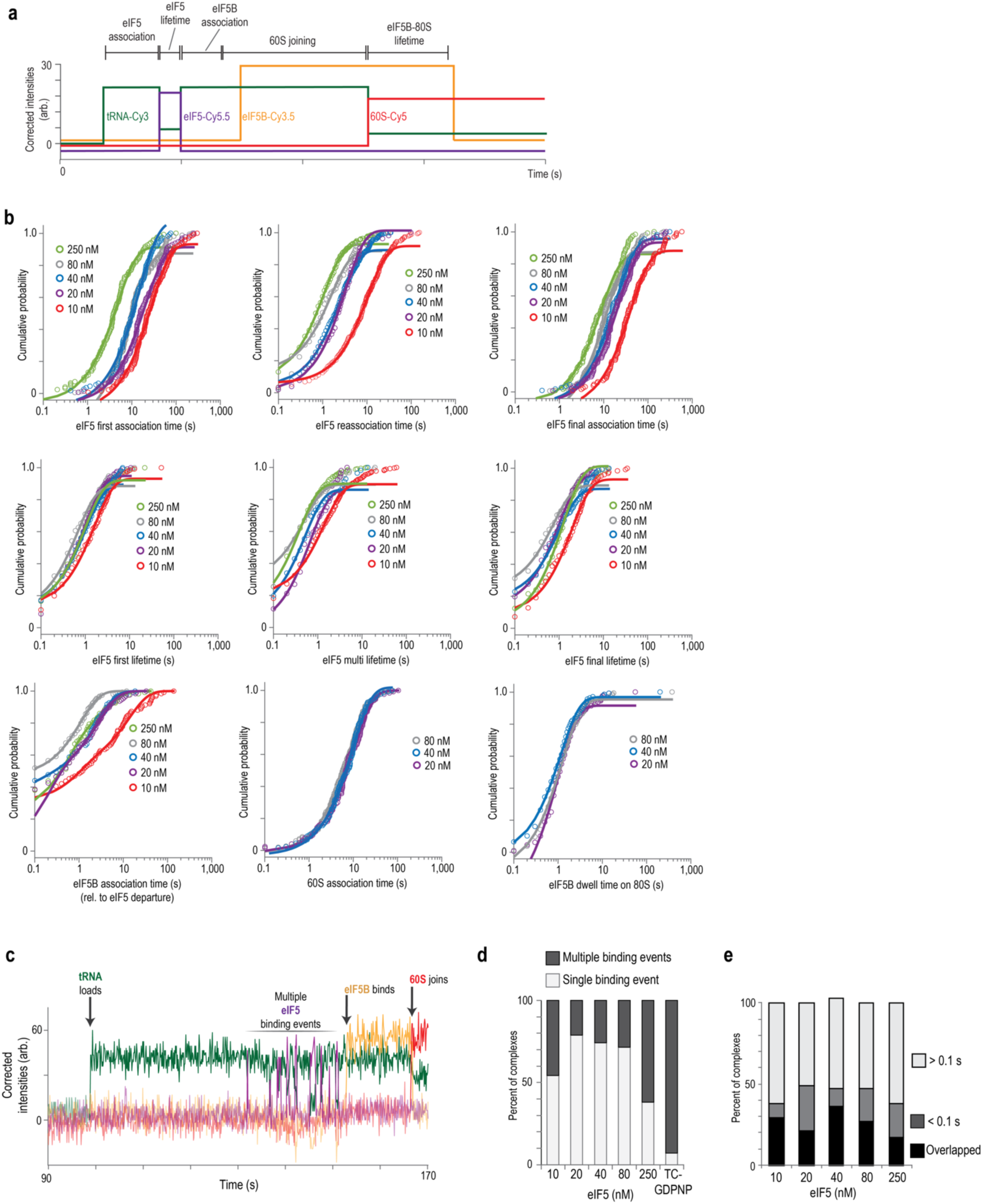
eIF5 transiently binds initiation complexes. (**a**) Schematic of predicted single-molecule data for real time eIF5-Cy5.5 experiments. Initial appearance of tRNA_i_-Cy3 (green) fluorescence indicates loading of the 43S initiation complex onto the mRNA. Either direct appearance of eIF5-Cy5.5 (purple) fluorescence or appearance of tRNA_i_-to-eIF5 FRET indicates the presence of eIF5 on initiation complexes. Appearance of eIF5B-Cy3.5 (orange) indicates association of eIF5B. Appearance of 60S-Cy5 (red) fluorescence indicates the 60S subunit joined to form the 80S ribosome; the relative proximity of tRNA_i_-Cy3 and 60S-Cy5 labeling sites in the 80S initiation complex yields low FRET. The kinetic parameters were defined as follows. ‘eIF5 first association time’ as the time elapsed from initial appearance of the tRNA_i_-Cy3 signal to initial appearance of eIF5-Cy5.5 signal. ‘eIF5 first lifetime’ as the duration of the first eIF5-Cy5.5 signal. ‘eIF5 reassociation time’ as the time elapsed from disappearance of the previous eIF5 signal to appearance of the next eIF5 signal. ‘eIF5 subsequent lifetimes’ as the duration of the subsequent eIF5 binding events. ‘eIF5 final association time’ as the time elapsed from appearance of the tRNA_i_-Cy3 signal until appearance of the final eIF5-Cy5.5 signal prior to eIF5B association. ‘eIF5 final lifetime’ was the duration of the final eIF5 binding event. Since most complexes contained a single eIF5 binding event, the ‘first’ and ‘final’ eIF5 binding events were identical on most complexes. ‘eIF5B association time’ was defined as the time elapsed from disappearance of the final eIF5 signal until appearance of the eIF5B-Cy3.5 signal. ‘60S association time’ was defined as the time elapsed from appearance of eIF5B-Cy3.5 signal until appearance of 60S-Cy5 signal. ‘eIF5B-80S lifetime’ was defined as the duration of the eIF5B-Cy3.5 signal on the 80S initiation complex. **(b)** Cumulative probability plots of the indicated kinetic parameters on β-globin mRNA at the indicated concentrations of eIF5-Cy5.5. **(c)** Example single-molecule data of an initiation complex that was bound multiple times by eIF5-Cy5.5. **(d)** Plot of the percent of initiation complexes that contained either a single or multiple eIF5-Cy5.5 binding events at the indicated eIF5-Cy5.5 concentrations. **(e)** Plot of the percent of initiation complexes where eIF5B associated either during (overlapped), within 100 ms (< 0.1 s), or after 0.1 s (> 0.1 s) of the final eIF5 binding event at the indicated concentrations of eIF5-Cy5.5. In all experiments, unlabeled eIF1 was present at a final concentration of 290 nM and eIF5-Cy5.5 was present at the indicated final concentrations (by Cy5.5 dye). See **Table S3** for the number of complexes and binding events analyzed in each experiment.

**Supplementary Figure S7.**
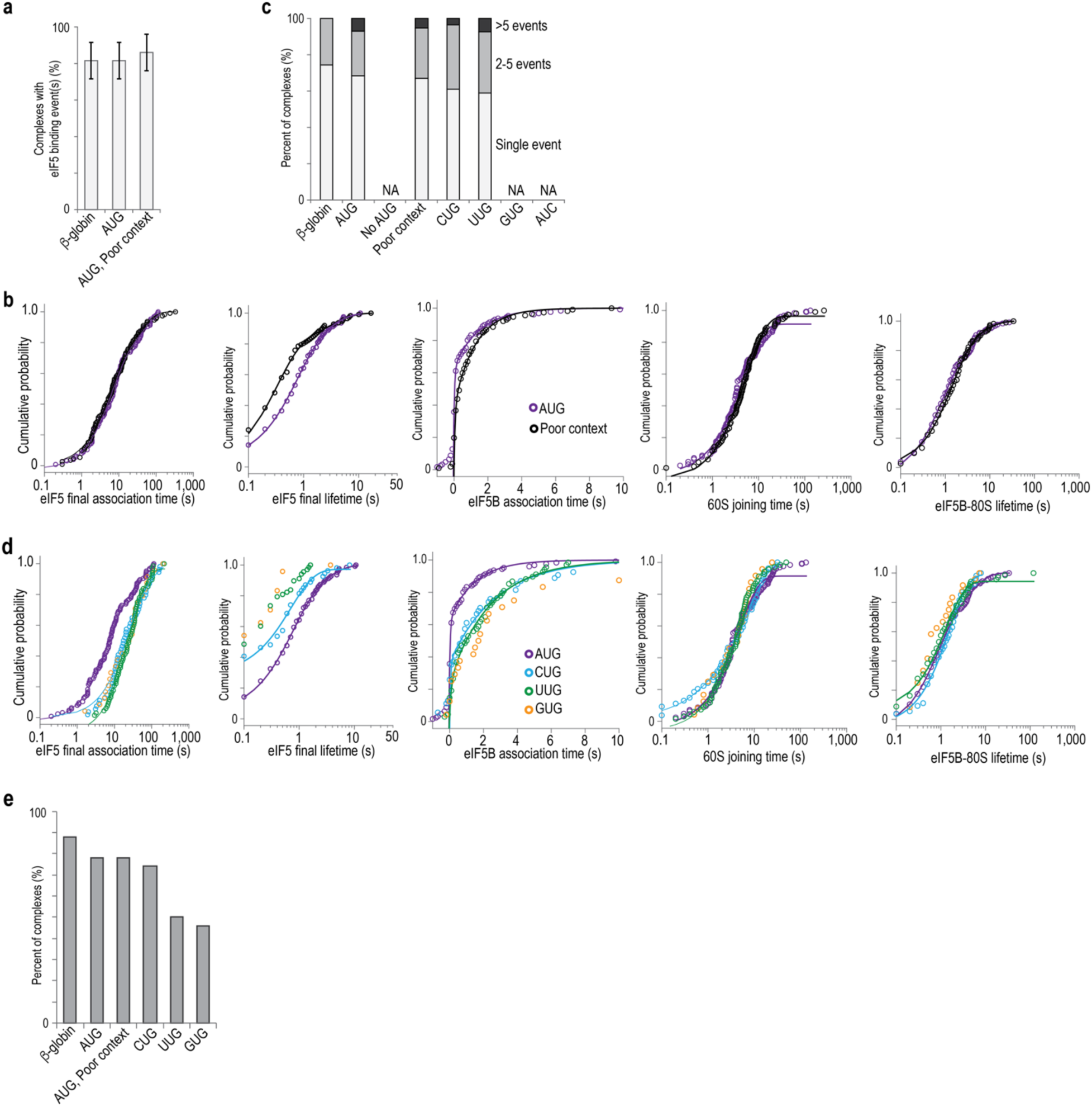
The identity of the translation start site alters eIF5 dynamics. (**a**) Plot of the percent of initiation complexes that contained at least one eIF5 binding event on the indicated mRNA. The percentage was corrected to account for an eIF5-Cy5.5 labeling efficiency of 50%. **(b,d)** Cumulative probability plots of the indicated kinetic parameters on the indicated mRNAs. **(c)** Plot of the percent of complexes that contained the indicated number of eIF5-Cy5.5 binding events on the indicated mRNAs. The percentage was corrected to account for an eIF5-Cy5.5 labeling efficiency of 50%. **(e)** Plot of the percent of initiation complexes that where the eIF5B association event either overlapped with or occurred within 100 ms of the final eIF5 binding event on the indicated mRNAs. In all experiments, unlabeled eIF1 was present at a final concentration of 290 nM and eIF5-Cy5.5 was present at 40 nM final concentrations (by Cy5.5 dye). See **Table S3** for the number of complexes and binding events analyzed in each experiment.

